# Transgenerational Epigenetic Inheritance Factors Localize to Spatially and Temporally Ordered Liquid Droplet Assemblages

**DOI:** 10.1101/220111

**Authors:** Gang Wan, Brandon D. Fields, George Spracklin, Carolyn Phillips, Scott Kennedy

## Abstract

Epigenetic information can be inherited for multiple generations (termed transgenerational epigenetic inheritance or TEI) ^1,2^. Non-coding RNAs have emerged as important mediators of TEI, although the mechanism(s) by which non-coding RNAs mediate TEI remains poorly understood. dsRNA-mediated gene silencing (RNAi) in *C. elegans* is a robust example of RNA-directed TEI^3–5^. To further our understanding of RNA-directed TEI, we conducted a genetic screen in *C. elegans* to identify genes required for RNAi inheritance. Our screen identified the conserved RNA helicase/Zn finger protein ZNFX-1 and the Argonaute protein WAGO-4. We find that WAGO-4 and ZNFX-1 act cooperatively in inheriting generations to maintain small interfering (si)RNA expression over generational time. ZNFX-1/ WAGO-4 localize to a liquid droplet organelle termed the P granule in early germline blastomeres. Later in development, ZNFX-1/WAGO-4 appear to separate from P granules to form independent foci that are adjacent to, yet remain distinct, from P granules. ZNFX-1/WAGO-4 labeled foci exhibit properties reminiscent of liquid droplets and we name these foci Z granules. In the adult germline, Z granules assemble into ordered tri-droplet assemblages with P granules and another germline droplet-like foci termed the *Mutator* foci. This work identifies a conserved RNA-binding protein that drives RNA-directed TEI in *C. elegans*, defines a new germline foci that we term the Z granule, demonstrates that liquid droplet formation is under developmental control, and shows that liquid droplets can assemble into spatially ordered multi-droplet structures. We speculate that temporal and spatial ordering of liquid droplets helps cells organize and coordinate the complex RNA processing pathways underlying gene regulatory systems, such as RNA-directed TEI.

---

Previous genetic screens have identified several components of the *C. elegans* RNAi inheritance machinery ^5–7^. We conducted a modified genetic screen designed to identify additional RNAi inheritance factors (Extended Data figure 1). Our screen identified 37 mutations that disrupted RNAi inheritance. We subjected DNA from these 37 mutant strains to whole-genome sequencing and identified four independent mutations in the gene *zk1067.2* (Fig. 1A). To confirm that *zk1067.2* is required for RNAi inheritance, we tested two additional alleles of *zk1067.2 (gk458570* and *gg561*) for RNAi inheritance defects. Treatment of *C. elegans* with *gfp* dsRNA (*gfp* RNAi) can trigger silencing of germline expressed *gfp* reporter genes for multiple generations ^3,5,7^. *gk458570* and *gg561* animals responded like wild-type animals to treatment with *gfp* dsRNA, however, progeny of these mutant animals were largely unable to inherit gene silencing (Fig. 1B, and Extended Data figure 2). Additionally, animals harboring *zk1067.2* mutations were defective for RNAi inheritance when the *dpy-11* and *oma-1* genes were targeted with dsRNA (Extended Data figure 3). We conclude that *zk1067.2* is required for RNAi inheritance.

**Figure 1.**
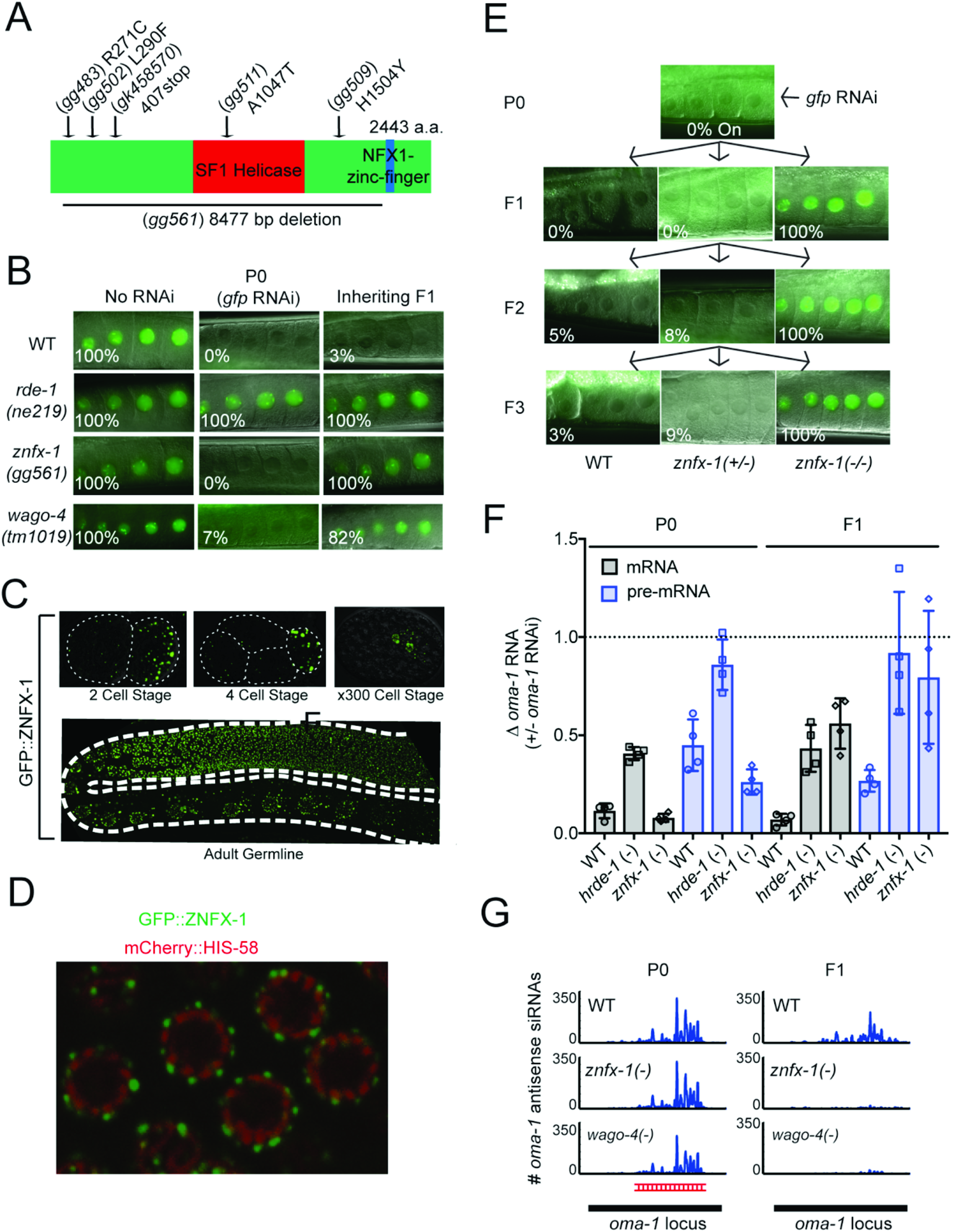
ZNFX-1 is a conserved RNA helicase required for RNAi inheritance in *C. elegans.* **A)** *znfx-1* alleles are indicated. **B)** Animals expressing a *pie-1∷gfp∷h2b* transgene ^7^ were exposed to *gfp* dsRNA. F1 progeny were grown in the absence of dsRNA, and GFP expression in oocytes was visualized by fluorescence microscopy. % of animals expressing *pie-1∷gfp∷h2b* is shown (n=3-6, at least 30 animals per condition and genotype were scored). P0, parental generation; WT, wild-type. Note, using a different *pie-1∷gfp∷h2b* reporter gene, we find that some *gfp* RNAi inheritance can be observed in *znfx-1* mutant animals (Fig. Extended Data Figure 2). **C)** Animals harboring a *gfp* epitope inserted into the *znfx-1* locus were visualized by fluorescence microscopy. **D)** Pachytene germ cells of animals expressing GFP∷ZNFX-1 and the chromatin marker mCherry∷HIS-58 ^33^ were visualized by fluorescence microscopy. **E)** *znfx-1(gg561)/+* heterozygous animals expressing *pie-1∷gfp∷h2b* transgene were exposed to *gfp* dsRNA. Progeny were grown in absence of *gfp* dsRNA for three generations. Images show GFP expression in oocytes. % of animals expressing *pie-1∷gfp∷h2b* is shown (n=30-65). **F)** WT, *hrde-1(tm1200)*, and *znfx-1(gg561)* animals were exposed to *oma-1* dsRNA. Total RNA from RNAi (P0) and inheriting (F1) generations was isolated, *oma-1* RNA levels were quantified using qRT-PCR using primers 5’ to the site of RNAi and data were normalized to *eft-3* pre-mRNA level. n=4. Error bars are +/− standard deviation of the mean (SD). Note, independent mRNA and pre-mRNA primer sets gave similar results (Extended Data figure 7). **G)** Secondary siRNA libraries (material and methods) were prepared from WT, *znfx-1 (gg561)*, and *wago-4(tm1019)* animals exposed to *oma-1* dsRNA (P0) and progeny of these animals (F1). Antisense reads mapping to *oma-1* locus are shown. Red indicates region of *oma-1* targeted by dsRNA. Reads counts were normalized to total number of sequenced reads (n=2).

Sequence analysis showed that ZK1067.2 encodes a 2443 amino acid protein that contains a superfamily one (SF1) RNA helicase domain and a Zn finger domain (Fig. 1A). A single putative ortholog of ZK1067.2 was found in most eukaryotic genomes. The *S. pombe* homolog of ZK1067.2 (termed HRR1) is a nuclear protein that physically interacts with Argonaute (AGO) and RNA Dependent RNA Polymerase (RdRP) to direct pericentromeric heterochromatin ^8^. The *N. crassa* homolog (SAD-3) is a cytoplasmic protein required for an RNAi-related phenomenon termed meiotic silencing of unpaired DNA (MSUD) ^9^. Homology between ZK1067.2 and its mammalian ortholog ZNFX1 extend to a Zn finger domain not present in fungal orthologues. We conclude that ZK1067.2 is a conserved protein that is involved in small RNA-mediated gene silencing in many eukaryotes. Henceforth, we refer to ZK1067.2 as ZNFX-1.

To begin to understand the function of ZNFX-1 during RNAi inheritance, we used CRISPR/Cas9 to insert a *3xflag∷gfp* epitope immediately upstream of the *znfx-1* start codon. *3xflag∷gfp∷znfx-1* animals behaved like wild-type animals during RNAi inheritance assays, indicating that *3xflag∷gfp∷znfx-1* encodes a functional protein (Extended Data figure 4). [Note, CRISPR-mediated gene conversion was used to epitope tag the genes described in this work and the resultant fusion proteins were expressed at or near wild-type levels and were functional for RNAi inheritance unless otherwise stated (also see Extended Data figure 4 and 5)]. We observed GFP∷ZNFX-1 expression in the adult germline as well as in developing germ cells during all stages of embryonic and larval development (Fig. 1C, and data not shown). No GFP∷ZNFX-1 expression was observed in somatic tissues. Post fertilization, *C. elegans* zygotes undergo a series of asymmetric cell divisions in which germline determinants segregate with the germline blastomeres of the P lineage. During embryonic development, ZNFX-1 foci were concentrated in, and segregated with, the germline P blastomeres (Fig 1C). In adult germ cells, GFP∷ZNFX-1 was concentrated in foci that were distributed in a perinuclear pattern around germ cell nuclei (Fig 1D). We conclude that *znfx-1* encodes a germline-expressed protein that segregates with the germline and localizes to perinuclear foci in adult germ cells.

ZNFX-1 could act in the parental generation to initiate RNAi inheritance or in progeny to receive RNAi inheritance signals (or both). The following results demonstrate that ZNFX-1 acts in inheriting generations to promote the memory of RNAi. We initiated gene silencing in *znfx-1/+* heterozygous animals and scored the +/+ and −/− progeny for their ability to inherit gene silencing. Progeny harboring at least one wild-type copy of *znfx-1* were capable of inheriting gene silencing while −/−progeny were not (Fig. 1E). When we initiated gene silencing in *znfx-1’(−/−)* animals, introduced a wild-type copy of *znfx-1* to progeny (via mating), and scored *znfx-1/+* cross-progeny for inheritance, we found that *znfx-1 (+/−)* progeny were able to inherit silencing (Extended Data figure 6). These data suggest that *znfx-1* acts in inheriting generations to enable RNAi inheritance. The following molecular data support this idea. First, treatment of animals with *oma-1* dsRNA silences the *oma-1* mRNA for multiple generations ^4,5^. We used qRT-PCR to measure *oma-1* mRNA and pre-mRNA levels in *znfx-1 (-)* animals exposed to *oma-1* RNAi, as well as in the progeny of these animals. *znfx-1(-)* animals responded normally to *oma-1* RNAi; however, their progeny failed to inherit silencing (Fig. 1F and Extended Data figure 7). Second, the nuclear RNAi factor NRDE-2 binds to pre-mRNA of genes undergoing heritable silencing ^5^. When *znfx-1(-)* animals were exposed directly to *oma-1* dsRNA, NRDE-2 bound to the *oma-1* pre-mRNA at wild-type levels, however, in progeny of *znfx-1(-)* mutant animals NRDE-2 failed to bind *oma-1* pre-mRNA (Extended Data Figure 8). Third, during RNAi inheritance, siRNAs with sequence complementarity to genes undergoing RNAi silencing are expressed for multiple generations ^5^. In *znfx-1(-)* animals exposed directly to *oma-1* dsRNA, *oma-1* siRNAs were produced at wild-type levels; however, the progeny of *znfx-1(-)* animals failed to express *oma-1* siRNAs (Fig. 1G). Together these data establish that ZNFX-1 is a dedicated RNAi inheritance factor.

The *C. elegans* genome encodes ~27 Argonaute (AGO) proteins. The molecular function of most of these AGOs remains a mystery. Two of the mutant strains identified by our genetic screen harbored mutations in the AGO-encoding gene *wago-4.* To confirm WAGO-4 is required for RNAi inheritance, we tested two additional *wago-4* deletion alleles (*tm1019* and *tm2401*) for RNAi inheritance defects. Both alleles exhibited RNAi inheritance defects (Fig. 1B and Extended Data Figure 9). Thus, like ZNFX-1, WAGO-4 is required for RNAi inheritance. Additionally, when we appended a *gfp* tag to the *wago-4* locus, we observed that, like ZNFX-1, WAGO-4 is a germline-expressed protein that segregates with the P lineage blastomeres and localizes to perinuclear foci (Extended Data Figure 9). For unknown reasons, our GFP∷WAGO-4 fusion protein was fully functional for RNAi inheritance in some RNAi inheritance assays but only partially functional in other assays (Extended Data Figure 10). TagRFP∷ZNFX-1 and GFP∷WAGO-4 colocalized in germ cells, hinting that WAGO-4 and ZNFX-1 may act together to promote RNAi inheritance (Fig. 2A). Three additional lines of evidence support this idea. First, consistent with the idea that *wago-4* and *znfx-1* are part of the same genetic pathway, a similar amount of residual RNAi inheritance was observed in both single and double *wago-4; znfx-1* mutant animals (Fig. 2B). Second, 3xFI_AG∷WAGO-4 co-precipitates with HA∷ZNFX-1, but not a HA tagged negative control protein HA∷HRDE-1, suggesting a physical interaction between the two proteins (Fig. 2C and Extended Data figure 11). Third, *wago-4* mutant animals behaved like *znfx-1* mutant animals in molecular assays of RNAi inheritance. For instance, following *oma-1* RNAi, *wago-4* mutant animals express wild-type levels of *oma-1* siRNAs; however, progeny of *wago-4* mutant animals fail to inherit these small RNAs (Fig. 1G). Taken together these data show that WAGO-4 functions with ZNFX-1 to transmit RNA-based epigenetic information across generations.

**Figure 2.**
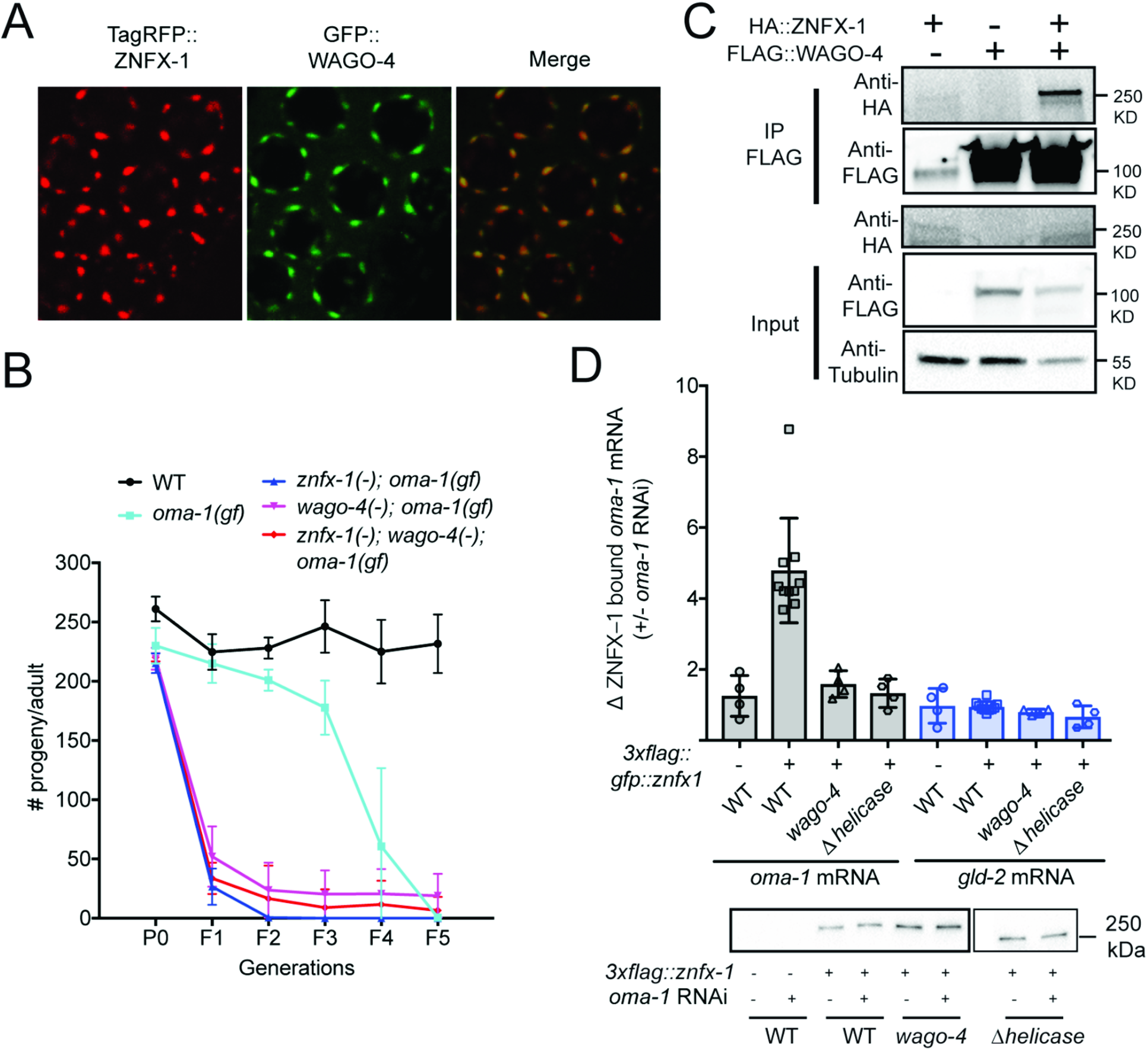
ZNFX-1 and WAGO-4 act cooperatively to drive RNAi inheritance. **A)** Fluorescent micrographs of pachytene germ cells of animals expressing GFP∷WAGO-4 and TagRFP∷ZNFX-1. **B)** *oma-1 (zu405) (gf)* is a gain of function temperature sensitive allele of *oma-1. oma-1(gf)* animals die at 20C unless *oma-1(gf)* is silenced by *oma-1* RNAi, which is heritable ^4^. *znfx-1(gg561), wago-4(tm1019)*, and double mutant animals are similarly defective for *oma-1(gf)* RNAi inheritance (n=3. +/− SD). **C)** ColP analysis of animals expressing *ha* or *3xflag* tags appended to *znfx-1* or *wago-4* loci, respectively (n=2). KD (Kilodaltons). **D)** Top panel, wild-type or 3xFLAG∷ZNFX-1 expressing animals were treated with *oma-1* dsRNA. ZNFX-1 was IP’ed in RNAi generation with a-FLAG antibodies and co-precipitating RNA was subjected to qRT-PCR to quantify *oma-1* mRNA co-precipitating with ZNFX-1 in wild-type, or *wago-4(tm1019)* animals. ZNFX-1 A *helicase* indicates 3xFLAG∷ZNFX-1 containing a 1487 bp in frame deletion of helicase domain, *gld-2* is a germline expressed control mRNA. n=4-10. +/− SD. Bottom panel, Western blot detecting 3xFLAG∷ZNFX-1 from one RNA IP replicate shown in top panel. Two unrelated lanes were removed from image. Note, these data show that, although RNAi defects aren’t observed in *znfx-1* mutants until inheriting generations, ZNFX-1 can interact with TEl-related RNAs in the RNAi generation.

How do WAGO-4 and ZNFX-1 promote RNAi inheritance? The closest homolog of ZNFX-1 is the RNA helicase SMG-2/UPF1, which binds and marks mRNAs containing premature termination codons ^10^. We wondered if, by analogy, ZNFX-1 might bind and mark mRNAs encoded by genes undergoing heritable gene silencing. To test this idea, we subjected 3XFLAG∷ZNFX-1 expressing animals to *oma-1* RNAi, immunoprecipitated 3XFLAG∷ZNFX-1, and used qRT-PCR to ask if *oma-1* RNAi directed ZNFX-1 to interact with *oma-1* mRNA. Indeed, *oma-1* RNAi caused ZNFX-1 to co-precipitate with *oma-1* mRNA (Fig. 2D). The following three lines of evidence show that the interaction of ZNFX-1 with TEl-related RNAs is a sequence-specific event directed by the RNAi machinery-First, RNAi targeting the *lin-15b* gene caused ZNFX-1 to interact with the *lin-15b* mRNA, but not the *oma-1* mRNA (and vice versa), indicating that ZNFX-1/mRNA interactions are sequence-specific (Extended Data Figure 12). Second, most RNA helicases bind RNA via their helicase domains. Deletion of the ZNFX-1 helicase domain did not affect ZNFX-1 expression (not shown) but did prevent ZNFX-1 from interacting with *oma-1* mRNA (Fig. 2D). Third, AGO proteins (like WAGO-4) use small RNA cofactors to direct sequence-specific RNA interactions. In *wago-4* mutant animals, ZNFX-1 failed to interact with the *oma-1* mRNA in response to *oma-1* RNAi (Fig. 2D). We conclude that RNAi directs ZNFX-1 to interact with mRNAs undergoing heritable silencing and that the AGO protein WAGO-4 is required for this property of ZNFX-1.

P granules are liquid droplet organelles that, like ZNFX-1/WAGO-4 foci, segregate with the germline blastomeres (P_0_-P_4_) during embryonic development ^11,12^. The low-complexity domain containing protein PGL-1 marks P granules ^13^. Both GFP∷ZNFX-1 and GFP∷WAGO-4 colocalized with PGL-1 ∷TagRFP in P_r_P_3_ germline blastomeres, suggesting that ZNFX-1 and WAGO-4 are P granule factors (Fig. 3A). MEG-3 and MEG-4 are low-complexity domain proteins that are redundantly required for P granule formation in the embryonic P lineage of *C. elegans* (Fig. 3B)^14^. In *meg-3/4(-)* embryos, ZNFX-1/WAGO-4 foci failed to form in the germline blastomeres (Fig. 3B). Thus, in early P_1_-P_3_ germline blastomeres, ZNFX-1 and WAGO-4 localize to P granules.

**Figure 3.**
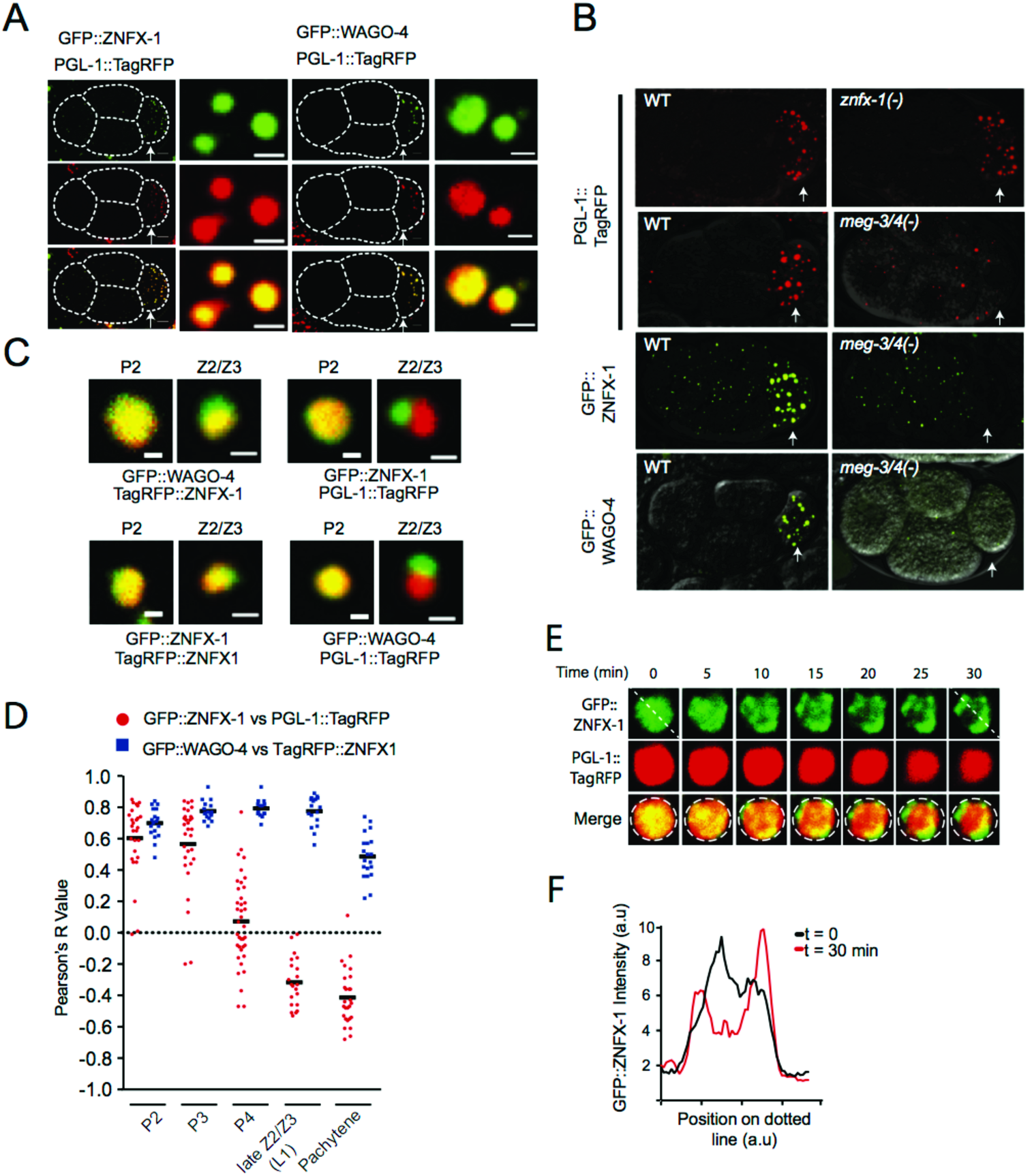
ZNFX-1/WAG0-4 appear to separate from P granules to form new foci during germline development. **A-B)** Micrographs of 4 cell embryos expressing indicated proteins are shown. P2 blastomeres are indicated by arrowheads. A) Magnifications of P granules are shown to the right. B) wild-type (WT), *znfx-1(gg561)*, or *meg-3(tm4259); meg-4(ax2026).* **C)** Magnifications of a P granule from a P2 blastomere and ZNFX-1/PGL-1 foci from Z2/Z3 cells from a larval stage one animal expressing indicated fluorescent proteins. **D)** Co-localization between GFP∷WAGO-4 and TagRFP∷ZNFX-1 (blue points) or GFP∷ZNFX-1 and PGL-1∷TagRFP (red points) was measured in individual granules at the indicated stages of development (see methods). Each data point represents an individual granule measurement (n = 15-35). **E)** Time-lapse micrographs of PGL-1∷TagRFP and GFP∷ZNFX-1 in early (-300 cell embryos) Z2/Z3 cells. (n>2). **F)** GFP intensity along dotted lines shown in **(E)**. Total amount of GFP∷ZNFX-1 fluorescence was relatively constant across time. Scale bars: **(A)** whole embryo 6 μn, individual granules 1 μm, **(C)** 0.5 μm.

At the ~100 cell stage of embryonic development the P_4_ blastomere divides to give rise to Z2 and Z3, which are the primordial germ cells of *C. elegans.* Surprisingly, we found that GFP∷ZNFX-1 no longer colocalized with PGL-1 ∷TagRFP in Z2/Z3 (Fig. 3C). Rather, GFP∷ZNFX-1 appeared in foci that were adjacent to (see below), yet distinct from, PGL-1 ∷TagRFP foci (Fig. 3C). Similar results were seen when antibodies were used to visualize PGL and ZNFX-1, indicating that failure to colocalize was not an artifact triggered by fluorescently conjugated epitopes (Extended Data figure 13). Quantitative analyses showed that the degree to which ZNFX-1 and PGL-1 colocalized changed during development, with a transition from colocalized to non-colocalized occurring between the P_3_ and Z2/Z3 stages (Fig. 3D). Conversely, GFP∷WAGO-4 and TagRFP∷ZNFX-1 remain colocalized throughout development (Fig. 3C/D). The ZNFX-1/WAGO-4 foci we see in Z2/Z3 could form *de novo* or by the separation of ZNFX-1/WAGO-4 and PGL-1 from within existing foci. We favor the later model as time-lapse imaging in early Z2/Z3 cells captured what appeared to be ZNFX-1 and PGL-1 separation events (Fig. 3E/F). The separation of ZNFX-1/WAGO-4 into discrete foci could be triggered by liquid liquid phase transitions or by the segregation of pre-existing states into discrete areas. We conclude that, late in germline development, ZNFX-1 and WAGO-4 are concentrated in foci that are adjacent to, yet are distinct, from P granules.

Liquid droplets organelles (such as the *C. elegans* P granules) are self-assembling cellular structures that form when specific proteins and RNAs undergo liquid-liquid phase transitions from surrounding cytoplasm. The ability of ZNFX-1 and WAGO-4 to separate from P granules hints that foci labeled by ZNFX-1/WAGO-4 (post Z2/Z3) may also be liquid droplets. Liquid droplets are typically spherical in shape and capable of flowing as distinct entities through other liquids, such as the cytoplasm. Consistent with the idea that ZNFX-1 foci are liquid droplets, we observed that during the course of oocyte maturation, spherically shaped ZNFX-1 foci transitioned from being perinuclear to being cytoplasmic, and as development continued, ZNFX-1 foci appeared farther and farther from nuclei, suggesting cytoplasmic flow (Fig. 4A). Constituents of liquid droplets undergo rapid internal rearrangements ^15,16^. Fluorescence recovery after photobleaching (FRAP) experiments showed that in ZNFX-1 foci, GFP∷ZNFX-1 fluorescence recovered from bleaching rapidly (t=~8 seconds), which is a rate similar to that reported for PGL-1 FRAP in P granules (Fig. 4B) ^11^. Taken together, these data show that ZNFX-1/WAGO-4 foci (post Z2/Z3) exhibit properties reminiscent of liquid droplets and, therefore, we refer to these foci as Z granules.

**Figure 4.**
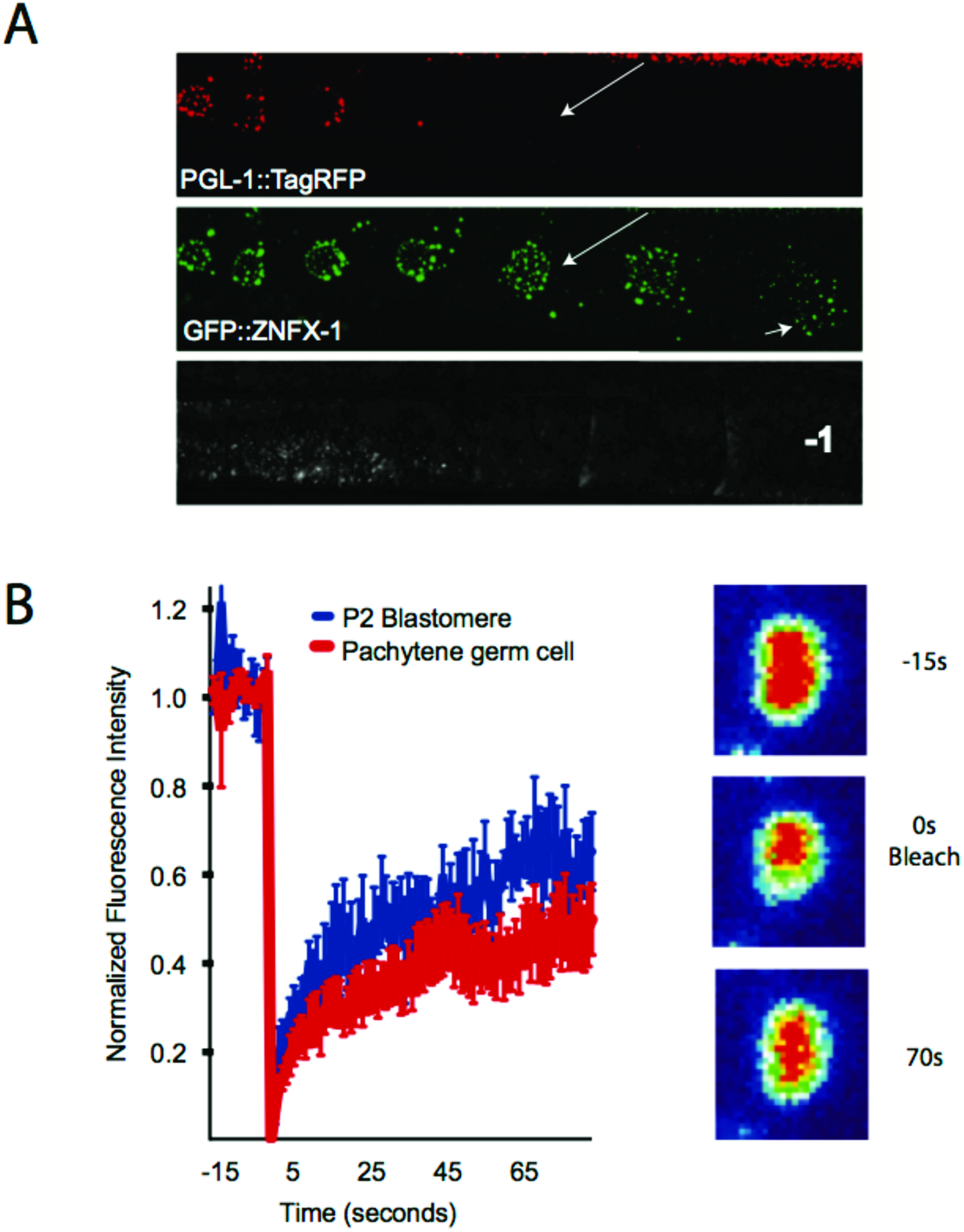
Z granules exhibit properties of liquid droplet organelles. **A)** Maturing oocytes of animals expressing the indicated fluorescent proteins. Long arrows indicates oocyte that contains Z granule, but not P granule. Short arrow indicates Z granules that are smaller and more distant from nuclei that Z granule at earlier developmental time points. −1 indicates −1 oocyte. **B)** GFP∷ZNFX-1 expressing animals were subjected to FRAP (see methods) and GFP fluorescence was monitored in bleached area over indicated times. Data is normalized to a non-bleached control granule from same sample, n = 7 granules +/− SEM. Right, heat maps (red represents high GFP signal) showing recovery of ZNFX-1 fluorescence in a representative bleached Z granule.

*C. elegans* germ cells possess at least two other foci (in addition to P granules) with properties akin to liquid droplets (processing bodies and *Mutator* foci) ^17,18^. TagRFP∷ZNFX-1 did not co-localize with MUT-16∷GFP, which marks *Mutator* foci, nor did GFP∷ZNFX-1 colocalize with mCherry∷PATR-1 or mRuby∷DCAP-1, which marks processing bodies (Fig. 5A and Extended Data Figure 14). Interestingly, although Z granules did not colocalize with *Mutator* foci the locations of these two foci were not random. Z granules were usually [89% of the time, (n=35)] found closely apposed to (no empty space observable between fluorescence signals) a *Mutator* foci (Fig. 5A). Similarly, Z granules were usually [91% of the time, (n=35)] found closely apposed to a P granule (Fig. 5A). EGO-1 is another marker of P granules ^17,19^. Z granules localized adjacent to, yet were distinct from, EGO-1 foci (Extended Data figure 15). Quantification of distances between surfaces and centers of fluorescence for the three foci supported the idea that Z granules localize adjacent to P granules and *Mutator* foci in adult germ cells (Fig. 5B). This analysis also showed that the distance between the surfaces of Z granules and P granules/Mi/tefor foci (but not P granules and *Mutator* foci) lies within the diffraction limit of light (zero distance values), indicating that Z granules exist in very close proximity to, and may be in direct physical contact with P granules and *Mutator* foci (Fig. 5B). Note, although Z granules are intimately associated with P granules/Mi/tefor foci in adult germ cells (and throughout most of germline development), they can exist independently. For instance, in the adult germline, P granules largely disappear during oocyte maturation (Fig.4A). Z granules remained visible at developmental time points when P granules were no longer present (Fig. 4A). Similarly, Z granules are present in developing germ cells at time points (*i.e.* P blastomeres) when *Mutator* foci are not thought to be present^17^. Shearing forces cause P granules in pachytene stage germ cells to disengage from nuclei and flow through the germline syncytium ^11^. After applying shearing force, we found that P granules flowed through the cytoplasm, however, Z granules remained static (Extended Data figure 16). Thus, Z granules can be separated from P granules and *Mutator* foci developmentally and physically. We conclude that Z granules represent an independent form of liquid droplet, which closely mirror P granules and *Mutator* foci in adult germ cells.

**Figure 5.**
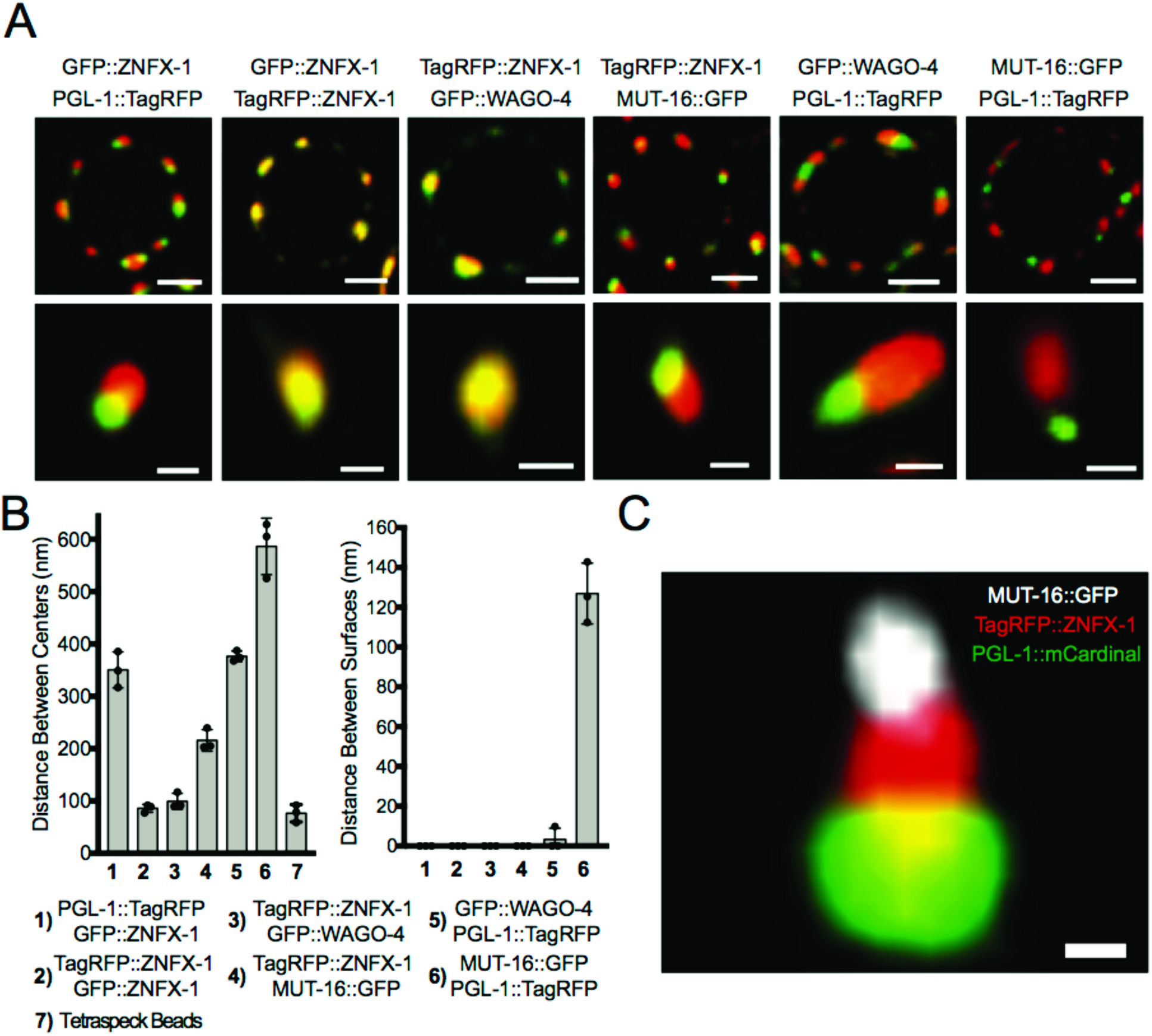
Z granules assemble into tri-droplet (PZM) structures with P granules and *Mutator* foci. **A)** Fluorescent micrographs of a single pachytene germ cell nucleus from animals expressing the indicated fluorescent proteins. 3D renders of representative foci are shown below. **B)** Distance between the centers (left) and surfaces (right) of the spaces occupied by the indicated fluorescent proteins was calculated as described in Methods. n=3 (10 granules per sample) +/− SD. Column 7 shows chromatic shift associated with imaging individual tetraspeck beads (zero distance). Data in right panel have been corrected for shift. **C)** Fluorescent micrograph from pachytene region germ cell. Image is magnification of a single tri-droplet assembly (PZM). Green=PGL-1∷mCardinal, Red=TagRFP∷ZNFX-1, and white=MUT-16 ∷GFP. Scale bars: **(A)** germ cell, 2 μm, single granule, 0.5 μm. **(C)** 0.25 μm. Position of nuclear membrane/nuclear pores with respect to each Z and M segments of PZM is not known (see Extended Data figure 21).

Our data suggest that Z granules may localize between (bridge) P granules and *Mutator* foci. To test this idea, we imaged the three foci simultaneously using PGL-1∷mCardinal, TagRFP∷ZNFX-1, and MUT-16∷GFP expressing animals ^20^. This analysis confirmed the idea that Z granules bridge P granules and *Mutator* foci (Fig. 5C): In 60% (52/86) of cases, we observed a Z granule in close apposition to both a P granule and a Mutator foci, while in 92% (48/52) of these cases the Z granule lay between the other two foci. In no case (0/52) did a P granule or a *Mutator* foci bridge the other two types of foci, respectively. Quantification of the distances between the centers and surfaces of Z granules, P granules, and *Mutator* foci from triple-marked images support the idea that Z granules act as a bridge between P granules and Mutator foci in adult germ cells (Extended Data figure 17). We conclude that P granules, Z granules, and *Mutator* foci form tri-droplet assemblages (henceforth PZMs) in adult germ cells and that the relative position of the three liquid droplets constituting the PZM is ordered.

Is PZM assembly required for RNA-directed TEI? Factors concentrated in Mutator foci^7,17,21,22^ and Z granules (this work) contribute to RNA-directed TEI. Additionally, we find that several factors, known to be required for P granule assembly, are needed for efficient RNAi inheritance (Extended Data figure 18). Thus, factors associated with all three segments of the PZM have now been linked to TEI. Additionally, we find that in animals with defective P granules, Z granules become malformed and ZNFX-1 fails to bind TEl-related RNAs, hinting that the segments of the PZM may communicate with each other during TEI (Extended Data figure 19). These results are consistent with the idea that PZM assembly contributes to RNA-directed TEI.

Here we show that the inheritance factors ZNFX-1 and WAGO-4 localize to a liquid droplet organelle that we name the Z granule. Given that Z granules segregate with the *C. elegans* germline, we speculate that one function of the Z granule is to concentrate and segregate silencing factors into the germline to promote RNA-based TEI. Consistent with this idea, we recently identified a mutation that inhibits the segregation of ZNFX-1 with the germline and showed that this same mutation is associated with RNAi inheritance defects (Extended Data Figure 20). ZNFX-1 is a conserved Zn finger RNA helicase, which, in *C. elegans*, marks RNAs produced by genes undergoing heritable silencing. The *S. pombe* ortholog of ZNFX-1 is Hrr1, which forms a nuclear complex (termed the RDRC) with Argonaute, the nucleotidyltransferase Cid12, and RdRP to amplify small RNA populations directing pericentromeric heterochromatin ^8^. We speculate that in *C. elegans* the RDRC has migrated from the nucleus to the Z granule where it promotes RNAi inheritance by: 1) binding inherited siRNAs (via WAGO-4), 2) marking mRNAs complementary to inherited siRNAs (via WAGO-4 and ZNFX-1), 3) using marked mRNAs as templates to amplify siRNA populations (via recruitment of one of the four *C. elegans* RdRP enzymes), and 4) repeating this cycle each generation (Extended Data figure 21). Support for this model comes from the recent identification of a homologue of the *S. pombe* RDRC factor CID12 (PUP-1/CDE-1) as a *C. elegans* RNAi inheritance factor^6^. ZNFX-1 and WAGO-4 appear to separate from components of the P granule during early embryogenesis to form independent liquid droplets. Separation occurs at a developmental time that correlates roughly with the first association of P granules with nuclear pores and the advent of germline transcription ^12,24–27^. We speculate that droplet separation might be triggered when newly synthesized mRNAs transit P granules and interact with RNA binding proteins to alter local protein concentration and initiate separation. In addition to temporal ordering, we find that Z granules are spatially ordered relative to P granules and *Mutator* foci, with Z granules forming the centerpiece of PZM tri-droplet assemblages in adult germ cells. These results show that cells possess mechanism(s) to organize and arrange liquid droplets in space as well as time. Additional work is needed to understand how PZM segments assemble in the correct order and if/how ordered PZM assembly contributes to RNA-based TEI. Small RNA-mediated gene regulatory pathways in animals are highly complex with thousands of small RNAs (e.g. miRNAs, piRNAs, siRNAs) regulating thousands of mRNAs at virtually all levels of gene expression. We speculate that the ordering of liquid droplet organelles in space and time helps organize and coordinate the complex RNA processing steps that underlie RNA-directed TEI (Extended Data figure 21). Similar strategies may be used by cells to organize and coordinate other gene regulatory pathways.

## Materials and Methods

### Strain list

N2 (CGC); (YY009) *eri-1(mg366)*, (YY193) *eri-1(mg366); nrde-2(gg91)*, (YY502) *nrde-2(gg91)*, (YY503) *nrde-2(gg90)*, (YY538) *hrde-1(tm1200)*, (YY562) *hrde-1(tm1200); oma-1(zu405)*, (YY913) *nrde-2(gg518[nrde-2∷3xflag∷ha])*, (YY916) *znfx-1(gg544[3xflag∷gfp∷znfx-1])*, (YY947) *hrde-1(tm1200); nrde-2(gg518)*, (YY967) *pgl-1(gg547[ pgl-1∷3xflag∷tagrfp])*, (YY968) *znfx-1(gg544); pgl-1(gg547)*, (YY996) *znfx-1(gg561)*, (TX20) *oma-1(zu405)*, (YY998) *znfx-1(gg544); ego-1(gg644[ha∷tagrfp∷ego-1])*, (YY1020) *znfx-1(gg561); oma-1(zu405)*, (SX461) *mjIS31(pie-1∷gfp∷h2b)*, (SS579) *pgl-1(bn101)*, (JH3225) *meg-3(tm4259); meg-4(ax2026)*, (DG3226) *deps-1(bn124)*, (YY1006) *eri-1(mg366); znfx-1(gg561)*, (YY1003) *eri-1(mg366); znfx-1(gk458570)*, (YY1021) *znfx-1(gg561); nrde-2(gg518)*, (YY1062) zn/x-7 (*gk458570)*, (YY1081) *deps-1(bn124); mjIS31*, (YY1083) *wago-4(tm1019)*, (YY1084) *wago-4(tm2401)*, (YY1093) *wago-4(tm1019); mjIS31*, (YY1094) *wago-4(tm2401); mjIS31*, (YY1108) *znfx-1(gg561); mjlS31*, (YY1109) *mjIS31; dpy-10(cn64)*, (YY1110) *eri-1(mg366); wago-4(tm1019)*, (YY1111) *eri-1(mg366); wago-4(tm2401)*, (YY1153) *wago-4(tm1019), znfx-1(gg544),mjlS31*, (YY1287) *znfx-1(gg611[ha∷znfx-1])*, (YY1305) *meg-3(tm4259); meg-4(ax2026), znfx-1(gg544)*, (YY1308) *meg-3(tm4259); meg-4(ax2026), pgl-1(gg547)*, (YY1325) *wago-4(gg620[3xflag∷gfp∷wago-4])*, (YY1327) *pgl-1(gg547); wago-4(gg620)*, (YY1364) *meg-3(tm4259); meg-4{ax2026), wago-4{gg620)*, (YY1388) *wago-4(gg627[3xflag∷wago-4])*, (YY1393) *znfx-1(gg611); wago-4(gg627)*, (YY1403) *wago-4(tm1019); znfx-1(gg561)*, (YY1408) *znfx-1(gg561); mjlS31; dpy-10(cn64)*, (YY1416) *znfx1(gg544);* axls1488, (YY1419) *znfx-1(gg561); wago-4(tm1019); oma-1(zu405)*, (YY1442) *znfx1(gg544);* hjSi397, (YY1446) *znfx-1(gg634[ha∷tagrfp∷znfx-1])*, (cmp3) *mut-16∷gfp∷flag+loxP*, (YY1444) *znfx-1{gg634); mut-16[mut-16∷gfp∷flag+loxP\*, (YY1452) *znfx-1(gg544); Itls37(pie-1 ∷mcherry∷his58)*, (YY1453) *znfx-1(gg634); wago-4(gg620)*, (YY1486) *znfx1(gg631[3xflag∷gfp∷znfx-1 Δ helicase])*, (YY1491) *wago-4(gg620); oma-1(zu405)* (YY1492) *pgl-1(gg640[pgl-1∷3xflag∷mcardinaf]); mut-16[mut-16∷gfp∷flag+loxP]; znfx-1(gg634)*, (YY1503) *pgl-1(gg547); mut-16[mut-16∷gfp∷flag+loxP]*.

### CRISPR/Cas9

gRNAs were chosen using Ape according to following standards: first, PAM sites are in the context of GGNGG ^28^ or GNGG; second, GC content of 20 bp spacer sequence was 40% to 60%; third, high specificity according to crispr.mit.edu. All CRISPR was done using co-CRISPR strategy ^29^. Plasmids were purified with PureLink− HiPure Plasmid Kits (Thermofisher). For deletions: two gRNAs (20ng/μl) were co-injected into gonads with pDD162(50ng/μl), *unc-58* gRNA(20ng/μl), AF-JA-76 (20ng/μl) and 1X taq buffer. For 3xFLAG or HA epitope tagging, single strand oligos (4nM ultramer from IDT, purified by isopropanol precipitation) with 50bp homology regions were used as repair templates. gRNA (20ng/μl) and repair template (20ng/μl) were co-injected into gonads with pDD162(50ng/μl), *unc-58* gRNA(20ng/μl), AF-JA-76 (20ng/μl) and 1X taq buffer. For GFP, tagRFP or mCardinal tagging, repair templates contained homologous arms of 500bp to 1000bp and were cloned into pGEM-7zf(+). Sequences were confirmed by Sanger sequencing. Repair templates were amplified with PCR, gel purified and isopropanol precipitated. PCR product was heated at 95 °C for 5min and then immediately put on ice for at least 2min. Injection mix was prepared: pDD162(50ng/μl), *unc-58* gRNA (20ng/μl), AF-JA-76(20ng/μl), gRNAs close to N terminal or C terminal of the genes (20ng/μl), heated and cooled repair template (50ng/μl) and 1X standard taq buffer. Injected animals were maintained at 25°C, Unc animals were isolated 4 days later and grown at 20°C. Animals were screened for deletion or tagging by PCR.

### RNA IP

Animals were flash frozen in liquid nitrogen and stored at −80°C. Animals were resuspended in sonication buffer (20 mM Tris.HCl PH 7.5, 200mM NaCl, 2.5mM MgCI2, 10% glycerol, 0.5% NP-40, 80U/ml RNaseOUT, 1mM DTT and protease inhibitor cocktail without EDTA) and sonicated (30s on, 30s off, 20%-30% output for 2min on a Qsonica Q880R sonicator, repeat once). Lysates were clarified by centrifuging at 14000 rpm for 15min. Supernatants were precleared with protein A agarose beads and incubated with FI_AG-M2 agarose beads for 2-3 hours at 4°C. Beads were washed with RIP buffer (20 mM Tris.HCl PH 7.5, 200mM NaCl, 2.5mM MgCI2, 10% glycerol, 0.5% NP-40) 6x. Protein and associated RNAs were eluted with 100ug/ml 3xFLAG peptide. RNAs were treated with Turbo DNase I for 20 min at 37°C and then extracted with TRIzol reagent followed by precipitation with isopropanol.

### RT-qPCR

mRNA isolated from total RNA or from RNA IP experiments, was converted to cDNA using the iScript cDNA synthesis kit according to vendor’s instructions. The following primer sequence were used to quantify mRNA levels, *oma-1* mRNA:GCTTGAAGATATTGCATTCAACC (Forward primer); AACTGTTGAAATGGAGGTGC (Reverse primer), *oma-1* pre-mRNA: GTGCGTTGGCTAATTTCCTG (Forward primer); CTGAATCGCGCGAACTTG (Reverse primer). *gld-2* mRNA: ACGTGTAGAAAGGGCTGCAC (Forward primer); GTCGATGCAGATGATGATGG (Reverse primer), *gld-2* pre-mRNA: CCTTATTAATTTCAGAGCTGCTGTC (Forward primer); AAGACTAGCACACGCAATCG (Reverse primer). *eft-3* pre-mRNA: CCTGCAAGTTCAACGAGCTTA (Forward primer); TGAAAAACAAATTGGTACATAAAC (Reverse primer).

### Mrt assay

Each generation, 3-6 L4 animals were picked to a single plate and grown at 25°C, average brood sizes were calculated by counting the total number of progeny per plate.

### RNAi inheritance assays

*dpy-11* and *gfp* RNAi inheritance: Embryos were collected via hypochlorite treatment and placed onto HT115 bacteria expressing dsRNA against *dpy-11* or *gfp.* F1 embryos were collected by hypochlorite treatment from RNAi or control treated adults and placed onto non-RNAi plates. Worms were scored at late L4 (*dpy-11*) or early young adult (*gfp*) stages, *oma-1* RNAi inheritance: Experiments were done at 20°C. Embryos were collected via hypochlorite treatment and placed onto HT115 bacteria expressing dsRNA against *oma-1.* 6 F1 embryos were picked onto a single OP50 plate. From F2 to F6, 6 L4 animals were picked onto a single OP50 plate. For genetic pathway analysis, *wago-4(tm1019); oma-1(zu405), znfx-1(gg561); oma-1(zu405)* and *wago-4(tm1019); znfx-1(gg561); oma-1(zu405)* animals were obtained from the same cross, *oma-1* RNAi inheritance assay were performed as described above. *tm1019* is a 571 bp deletion which removes part of the PIWI domain. *tm1019* also introduces a frameshift that would be expected to prevent translation of the rest of the PIWI domain, *znfx-1 (gg561)* is a 8476 bp deletion that deletes most (2300/2400 a.a.) of ZNFX-1, including the helicase domain. Both alleles were presumably null.

### Co-immunoprecipitation

Young adults were flash frozen in liquid nitrogen. Animals were ground into powder in liquid nitrogen and resuspended in 1ml 1X lysis buffer (20mM Hepes pH7.5, 100mM NaCl, 5mM MgCI2, 1mM EDTA, 10% Glycerol, 0.25% Triton, 1mM fresh made PMSF, 1X complete protease inhibitor from Roche without EDTA) and rotated for 45min at 4°C. Lysate was cleared by spinning at 5000 rpm for 15min, 30μl protein G beads were added to preclear lysate for 30min. 3xFLAG∷WAGO-4 proteins were pulled down by 30μl agarose beads conjugated to a-FI_AG antibody (A2220, Sigma-Aldrich). Input and IP proteins were separated by SDS-PAGE and detected by FLAG M2 antibody and HA antibody (Roche, 3F10).

### Small RNA sequencing

Total RNA was extracted using TRIzol. 20ug total RNA were separated by 15% Urea gel. Small RNA from about 18 nt to 35 nt were cut from gel. Small RNAs were cloned using a 5’ monophosphate independent small RNA protocol as previously described ^30^. Libraries were multiplexed with a 4 nt 5’ barcode and a 6 nt 3’ barcode and pooled for next generation sequencing on a NextSeq 500. FastX 0.0.13 was used to separate reads that contained the 3’ adapter and filter low quality reads for further analysis. Reads >14 nt were mapped to the *C. elegans* genome (WS220) using Bowtie. Read counts were normalized to the total number of reads matching the genome. Two independent libraries were prepared and the two replicates were combined for Figure 1.

### Microscopy and Analysis

To image larval and adult stages, animals were immobilized in M9 with 0.1% Sodium Azide, and mounted on glass slides. To image embryos, gravid adults were dissected on a coverslip containing 10 μl of 1X egg buffer, and then mounted on freshly made 3% agarose pads. Animals were imaged immediately with a Nikon Eclipse Ti microscope equipped with a W1 Yokogawa Spinning disk with 50 μn pinhole disk and an Andor Zyla 4.2 Plus sCMOS monochrome camera. A 60X/1.4 Plan Apo Oil objective was used unless otherwise stated.

### Colocalization

The degree of colocalization between different fluorescently labeled proteins across development was calculated using the Coloc2 plugin from ImageJ. Animals were imaged as described above with the exception of using a 100X/1.45 Plan Apo Oil objective. 3-5 granules were selected from at least 3 different animals across each stage of development specified. Region of interest (ROI) masks were generated using the 3D ROI Manager plugin in ImageJ to eliminate black regions surrounding granules ^31^. Coloc2 was used to generate a Pearson’s R Value for degree of colocalization between two channels in the region defined by the ROI mask.

### FRAP

Fluorescence recovery after photobleaching (FRAP) experiments were conducted using a Zeiss LSM 780 point scanning confocal equipped with a Quasar PMT x2 + GAaSP 32 Channel Spectral Detector using a 63X/1.4 Plan Apo Oil objective. Adult animals (for pachytene germ cells) or embryos (for P2 blastomere) were suspended in a mixture of 0.5% sodium azide and 50% 0.1 μm polystyrene beads (Polysciences) to inhibit movement. The mixture was added to a coverslip and placed on a fresh 3% agarose pad. Slides were sealed with nail polish. The bleaching plugin within the Zeiss Black software was used to specify the ROI to be bleached. One ROI was used for all data points. Single z slice images were acquired at 1 second intervals for 15 seconds, followed by bleaching, then continued at 1 second intervals for 85 seconds. Images were aligned using neighboring granules in ImageJ to account for subtle shifts in movement. An ROI was generated around the bleached region and continuously measured across all time points using the plot profile function within ImageJ. Data was normalized to an unbleached control granule to account for background bleaching throughout the 100 second period. Normalized data points were averaged across all 7 granules and plotted using Prism. The heat map of a representative granule was generated using the thermal LUT within ImageJ.

### Quantification of distances between foci centers and surfaces

We imaged pachytene germ cell nuclei in 3 animals. ~10 granules were selected from each animal. Confocal z stacks were opened with the 3D objects counter plugin from ImageJ to generate x, y, and z coordinates for the center of each object^32^. To account for chromatic shift between channels, 0.1 μm Tetraspek beads were imaged and granule distances were corrected accordingly. Distances between foci surfaces was calculated with 3D ROI manager in ImageJ^31^. Thresholding function within 3D ROI manager was used to eliminate background signal.

### Immunofluorescence

~30 animals were sliced open in 8 ul of 1X egg buffer (25 mM HEPEs, pH 7.3, 118 mM NaCl_2_, 48 mM KCl, 2 mM CaCI_2_, 2 mM MgCl_2_) to isolate gonads and embryos. A coverslip was added and slides were placed on a metal block (chilled on dry ice) for 10 minutes. Coverslips were popped off and slides were submerged in methanol at −20°C for 10 minutes, followed by acetone at −20°C for 5 minutes. Samples were allowed to dry at room temperature for 3 minutes. 500 ul of 1XPBST was added to each sample and incubated for 5 minutes at room temperature followed by 500 ul of 1XPBST + 1% BSA for 30 minutes at room temperature. 50 ul of antibody solution (1XPBST, 1% BSA, 1:20 dilution of anti PGL-1 antibody (K76 from DSHB), and 1:250 dilution of anti HA antibody (abcam ab9110)) was added to each sample. Samples were covered with parafilm and incubated overnight at room temperature inside a humid chamber. Samples were washed 3 times in 1XPBST at room temperature for 10 minutes. Secondary antibodies (Alexa Fluor 555 goat anti-rabbit (Life technologies A21429) and Alexa Fluor 488 goat anti-mouse (Life technologies A10667) were diluted 1:50 in 1XPBST 50 ul of secondary solution was added to each sample, covered with parafilm, and incubated for 90 minutes in the dark at room temperature. Samples were washed 3 times in 1XPBST at room temperature for 10 minutes. 15 ul of Vectashield antifade + DAPI was added to each sample. Slides were sealed with nail polish.

---

**Extended Data figure 1.**
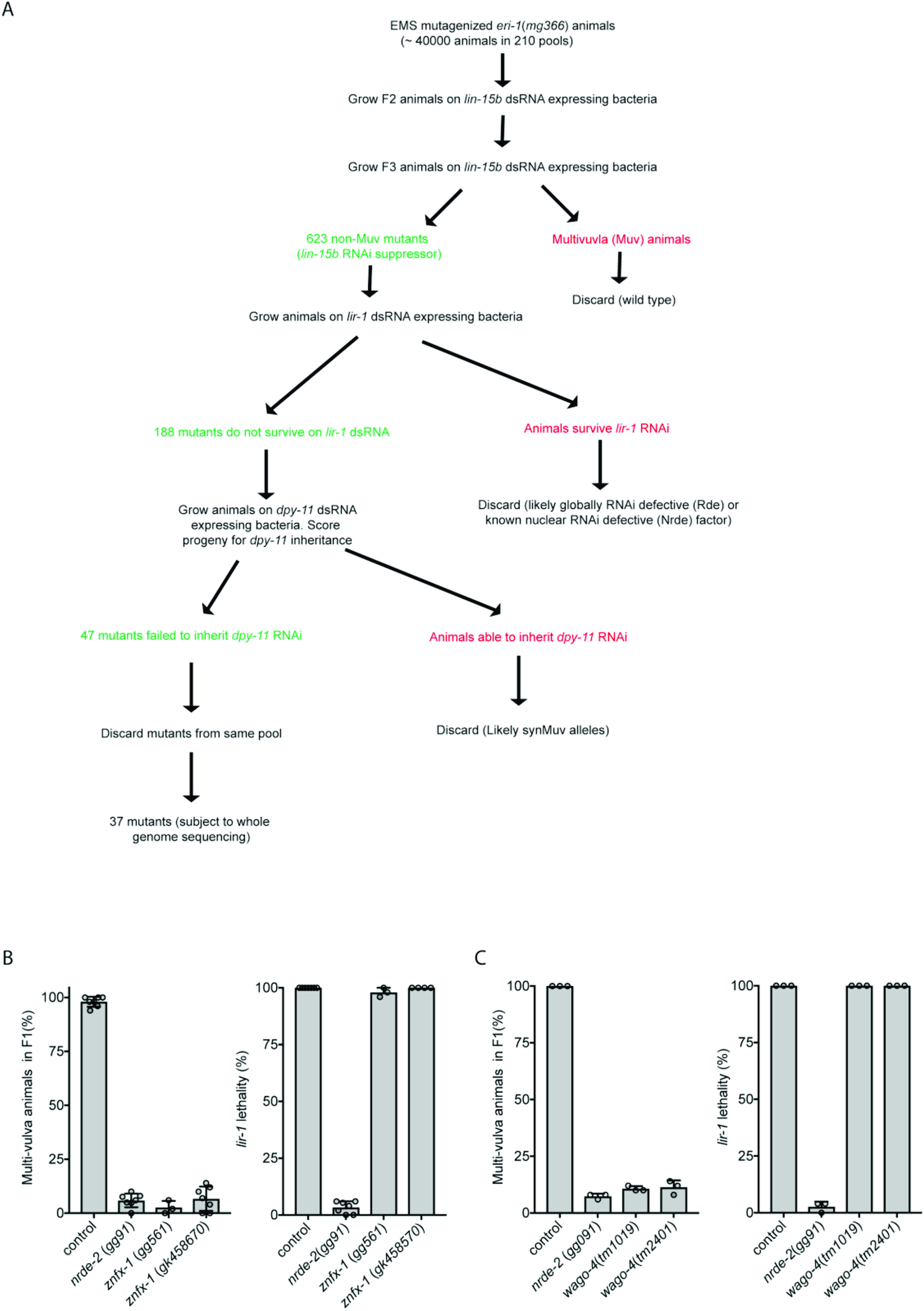
Genetic screen to identify novel RNAi inheritance mutants. **A)** We conducted a genetic screen to identify components of the *C. elegans* RNAi inheritance machinery. The screen contained several filters (see below) to remove known RNAi inheritance factor and, therefore, to allow us to focus on new RNAi inheritance factors. We know that factors defective for RNAi inheritance are also defective for nuclear RNAi (Buckley et al. 2012). Therefore, our screen began with selections for mutant alleles that disrupt nuclear RNAi. We have developed two selections for nuclear RNAi mutants. First, The *lin-15b* and *lin-15a* genes are transcribed as a polycistronic message that is spliced within the nucleus into *lin-15b* and *lin-15a* mRNAs (Blumenthal et al. 2002). Animals harboring mutations in both *lin-15b* and *lin-15a* exhibit a multivulva (Muv) phenotype (Clark, Lu, and Horvitz 1994; Huang, Tzou, and Sternberg 1994). RNAi targeting *lin-15b* (in *eri-1(-)* animals) silences *lin-15b* and *lin-15a* co-transcriptionally, thus inducing a Muv phenotype (Guang et al. 2008). The previously identified nuclear RNAi factors are required for *lin-15b* RNAi-induced co-transcriptional silencing of *lin-15a* and, therefore, for *lin-15b* RNAi induced Muv. A second assay for nuclear RNAi is *lir-1* RNAi. *lir-1* RNAi is lethal because *lir-1* is in an operon with *lin-26*, and co-transcriptional silencing of *lin-26* by *lir-1* RNAi causes lethality (Guang et al. 2008). Nuclear RNAi defective (NRDE) animals do not die in response to *lir-1* RNAi because they fail to silence *lin-26* (Guang et al. 2008). Previous genetic screens in the lab have used suppression of //’/"-1 RNAi to find factors required for nuclear RNAi. These screens have reached saturations: we have identified multiple alleles in all the *nrde* genes using this approach. Unpublished work from the lab shows, however, that hypomorphic alleles of the *nrde* genes will often block *lin-15b* RNAi-induced Muv and yet still die in response to *lir-1* RNAi. We interpret these data to mean that survival from *lir-1* RNAi is a much stronger selection for nuclear RNAi mutants than a failure to form Muv in response to *lin-15b* RNAi. In other words, factors contributing to nuclear RNAi, but not being 100% required for nuclear RNAi, would not be identified by *lir-1* RNAi suppression screens. For these reasons, our screen looked for suppressors of *lin-15b* RNAi, which did not suppress *lir-1* RNAi, because we felt this screen might identify genes missed in our previous genetic screens. **Step #1. Identify factors required for nuclear RNAi.** *eri-1(mg366)* animals were mutagenized with EMS. F2 progeny were exposed to bacteria expressing *lin-15b* dsRNA. Non-Muv animals were kept as candidate novel nuclear RNAi factors. **Step #2. Discard known nuclear RNAi factors.** We know all non-essential genes that can mutate to suppress *lir-1* RNAi. Therefore, we discarded mutants that suppressed *lir-1* RNAi as these alleles are likely known nuclear RNAi factors. Mutants that did not suppress *lir-1* may harbor mutations in factors important, but not essential, for nuclear RNAi. **Step #3. Identify mutations that suppress RNAi inheritance.** The last filter in our screen was to identify mutant alleles that disrupted RNAi inheritance. To do this, we subjected remaining mutant animals to *dpy-11* RNAi. *dpy-11* RNAi causes animals exposed to *dpy-11* dsRNA to become Dumpy (Dpy). Progeny of animals exposed to *dpy-11* dsRNA inherit *dpy-11* silencing and are Dpy (Burton, Burkhart, and Kennedy 2011). RNAi inheritance mutants become Dpy in response to *dpy-11* RNAi, however, the progeny of these animals fail to inherit *dpy-11* silencing, and, therefore, are not Dpy. Thus, any of our mutant animals that became Dpy in response to *dpy-11* RNAi, but whose progeny were not Dpy, were kept for further analysis. Finally, only one mutant was kept from each pool (pools were maintained as independent populations throughout the screen). B-C) The data in panels B and C show that the alleles of *znfx-1* and *wago-4* identified by our screen are (as expected) defective for *lin-15b* RNAi and not defective for *lir-1* RNAi (n=3-6, +/− SD).

**Extended Data figure 2.**
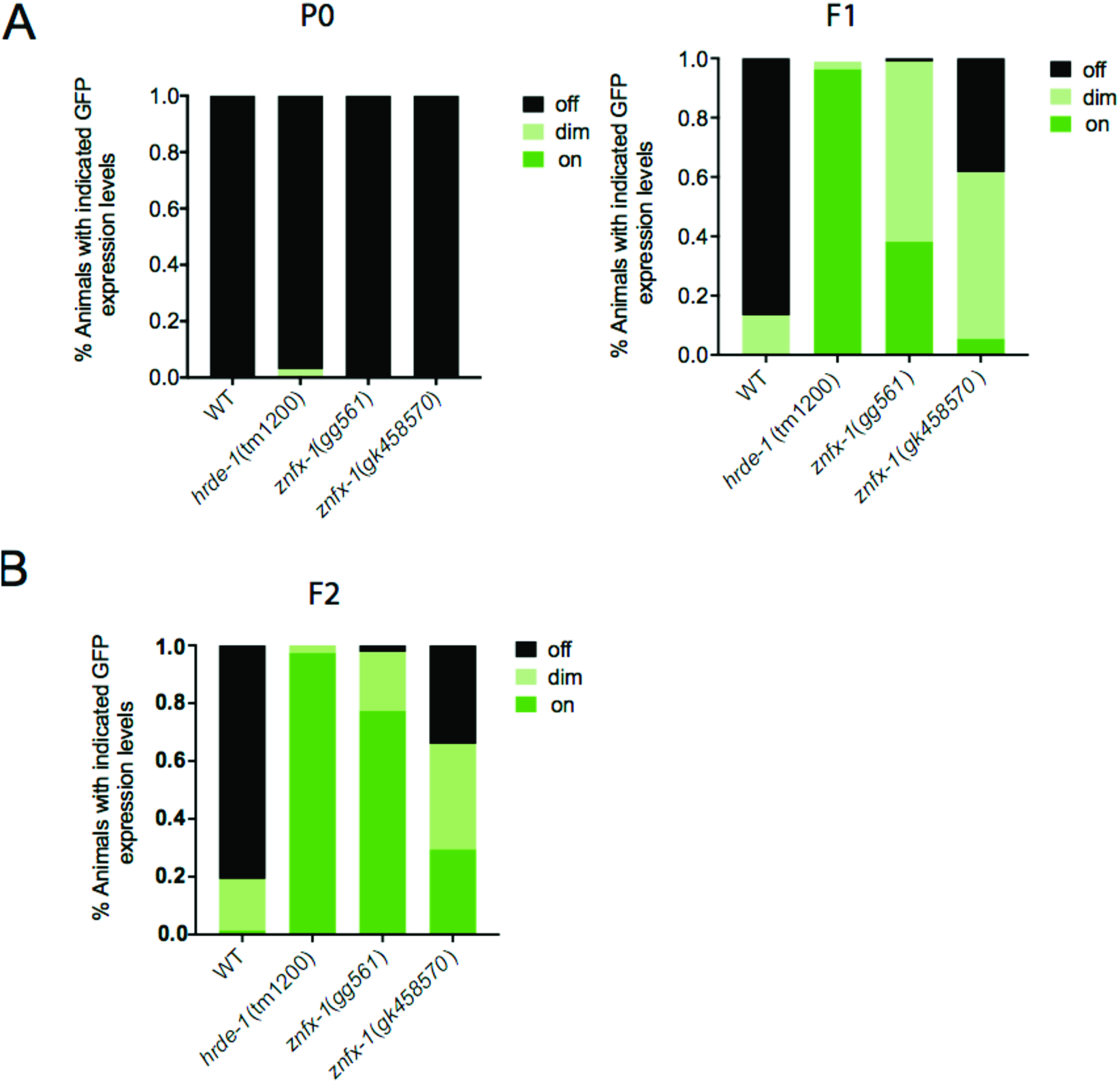
Confirmation that ZNFX-1 promotes transgenerational *gfp* RNAi inheritance. **A-B)** Animals expressing a *pie-1∷gfp∷h2b* transgene were exposed to *gfp* dsRNA (Vastenhouw et al. 2006). The % of P0, F1 and F2 progeny, of the indicated genotypes, expressing GFP was quantified. Data represent scoring of at least 80 animals in each generation and for each genotype. Note, the *gfp* reporter transgene used in this study is a multi-copy version of the single copy version used in Figure. 1b of the main text. Also note, that some RNAi inheritance can be seen in *znfx-1* mutant animals using this reporter transgene.

**Extended Data figure 3.**
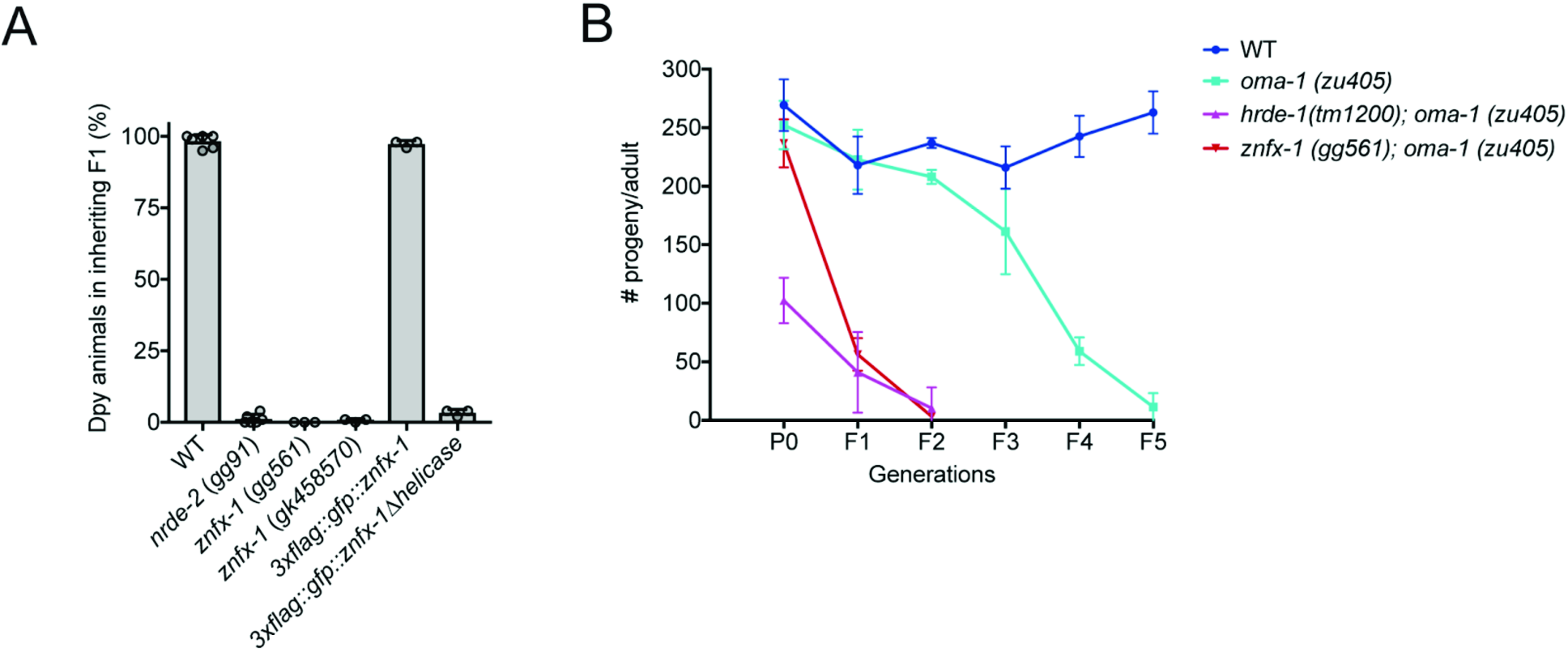
*znfx-1* is required for heritable silencing of endogenous genes. **A)** Animals of indicated genotypes were exposed to *dpy-11* dsRNA. The F1 progeny of these animals, were grown in the absence of *dpy-11* dsRNA, and were scored for Dpy phenotypes. (n>3, bars, S.D). Consistent with the idea that ZNFX-1 (and NRDE-2) is required specifically for inheritance, *znfx-1* mutant animals exposed directly to *dpy-11* dsRNA are Dpy (data not shown). **B)** *zu405ts* is a temperature sensitive (ts) lethal (embryonic arrest at 20°C) allele of *oma-1* (Lin 2003). *oma-1* RNAi suppresses *oma-1(zu405ts)* lethality, and this effect is heritable (Alcazar, Lin, and Fire 2008; Buckley et al. 2012). Animals of the indicated genotypes were exposed to *oma-1* dsRNA and the fertility of the progeny of these animals was scored over generations. (n=3, +/− SD). Data show that *znfx-1* mutant animals are defective for *oma-1* RNAi inheritance.

**Extended Data figure 4.**
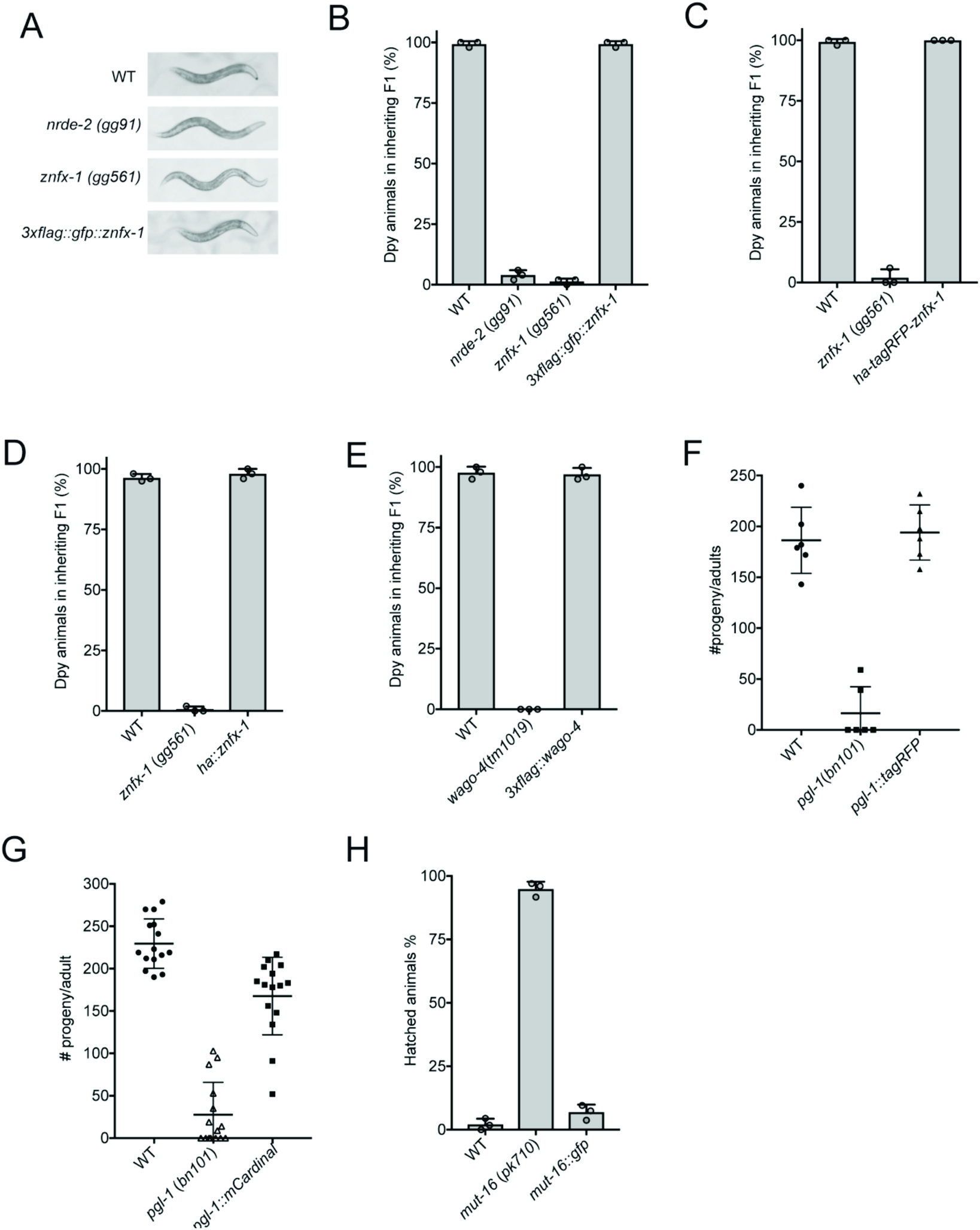
CRISPR/Cas9-epitope tagged genes used in this study produce functional proteins. Panels **(A-E)** show that the addition of epitope tags by CRISPR/Cas9-mediated gene conversion of *znfx-1* or *wago-4* did not affect function of ZNFX-1 or WAGO-4 in RNAi inheritance. Data shows results from *dpy-11* RNAi inheritance assay in which the progeny of animals exposed to *dpy-11* dsRNA are visually scored for the inheritance of Dpy phenotypes. (n=3, bars, S.D) **F-G)** *pgl-1* mutant animals show a temperature sensitive (25C) sterile phenotype. Panels **F** and **G** show that our addition of epitope tags (by CRISPR/Cas9-mediated gene conversion) to *pgl-1* locus did not affect PGL-1 function as these animals are fertile. L4 animals were singled from 20°C to 25°C and brood sizes were scored. *pgl-1-tagrfp* (n=6) and *pgl-1∷mcardinal; tagrfp∷znfx-1; mut-16∷gfp* (n=15). **H)** *mut-16(-)* animals are defective for *pos-1* RNAi. Embryos of indicated genotype were grown on *pos-1* dsRNA expressing bacteria. 6 L4 animals were picked to *pos-1* dsRNA expressing bacteria and laid egg overnight. Unhatched embryos and hatch animals were scored. The data shows that addition of *gfp* to *mut-16* locus did not affect MUT-16 function. (n=3, bars, S.D).

**Extended Data figure 5.**
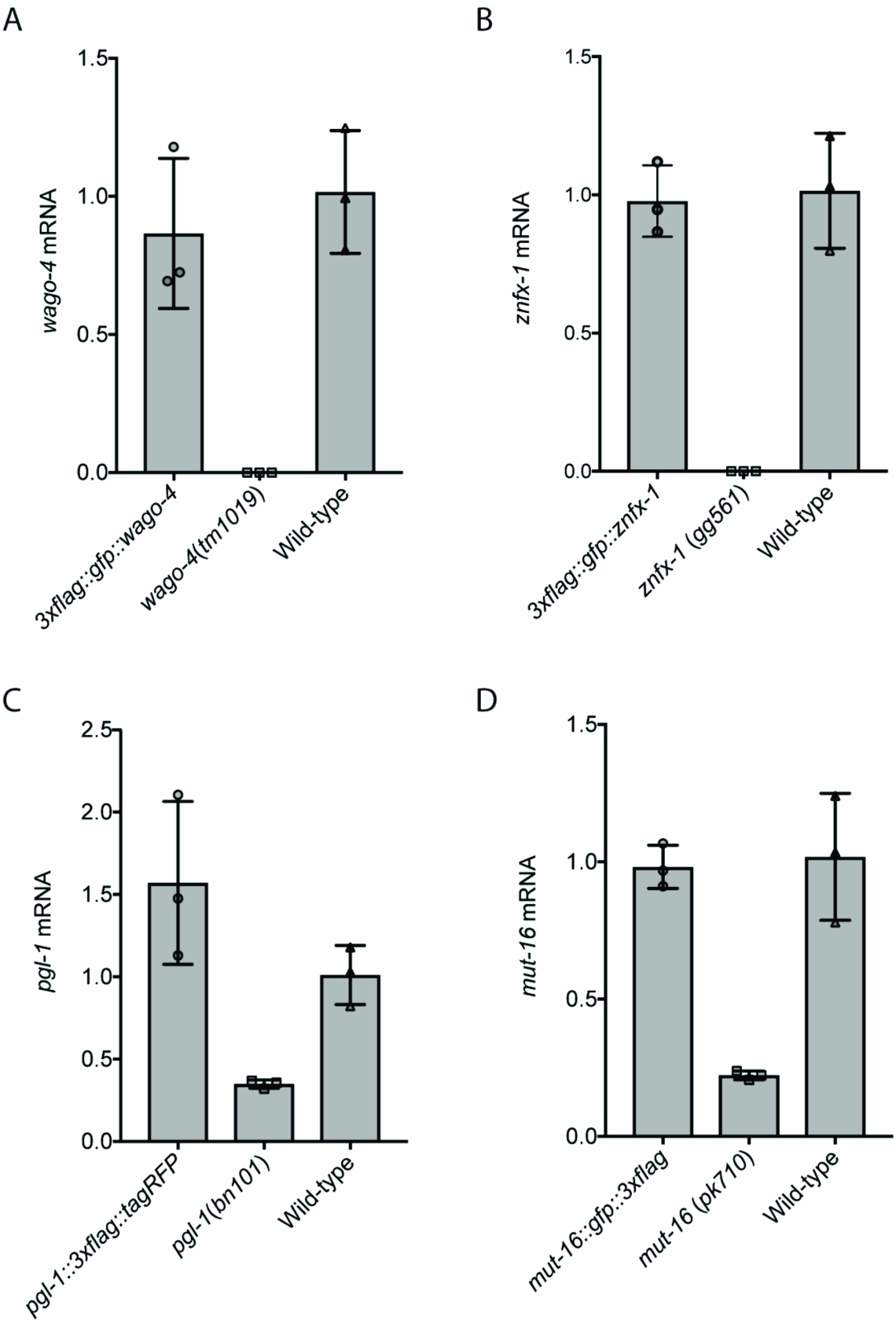
CRISPR/Cas9 epitope tagging of genes in this study did not substantially alter gene expression. **(A-D)** To address the possibility that epitope tagging of the genes used in this study changed gene expression levels we isolated total RNA from animals of the indicated genotypes and used qRT-PCR to quantify indicated mRNA levels. Primers target exon-intron junctions. Early stop or deletion alleles for each of these loci were used as controls. *wago-4(tm1019)* and *znfx-1(gg561)* are deletions and primers were located within deleted regions. *pgl-1(bn101)* and *mut-16 (pk710)* are nonsense alleles. Decrease of mRNA levels in these mutants is likely due to nonsense mediated decay (n=3, +/−SD).

**Extended Data figure 6.**
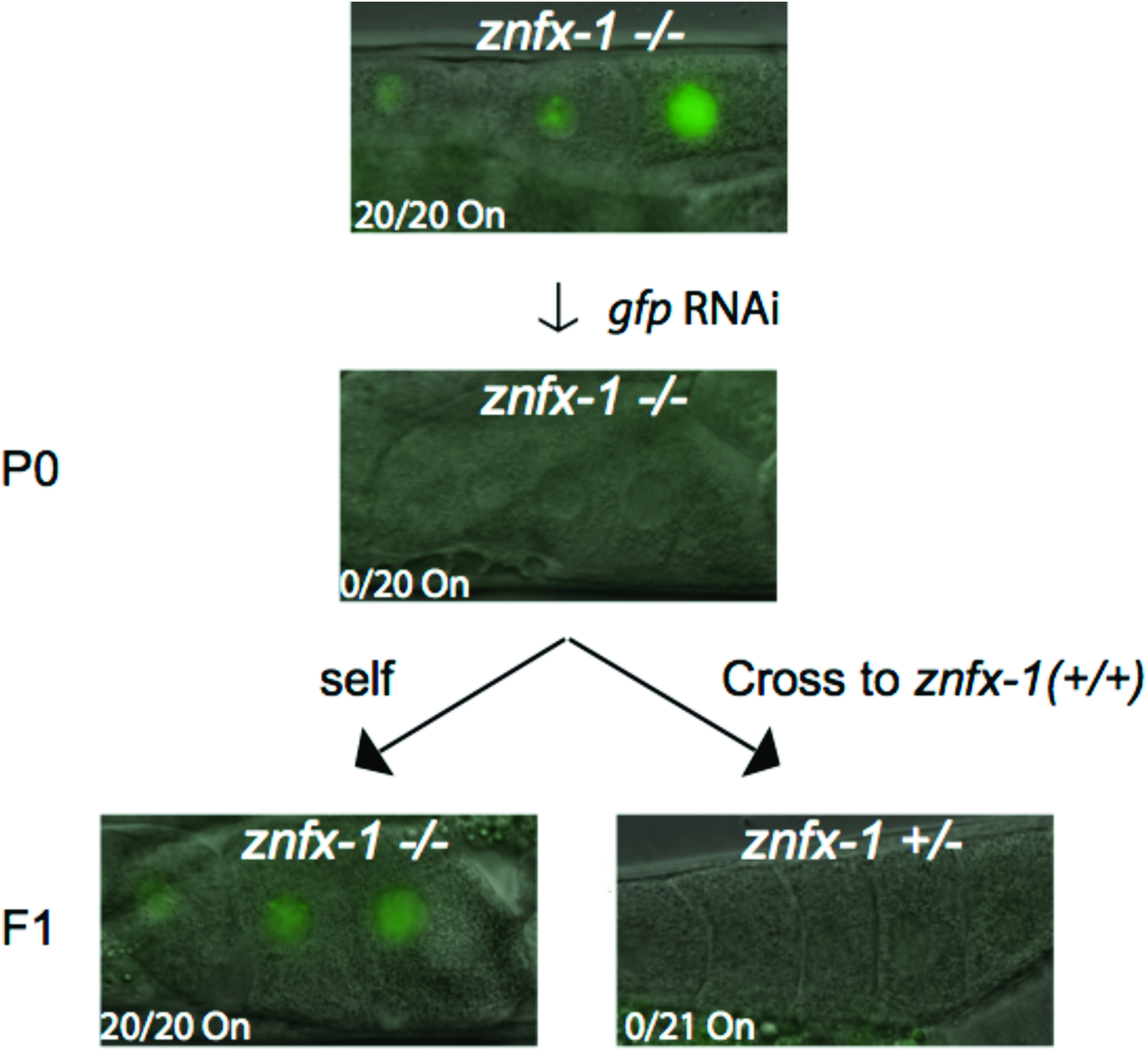
ZNFX-1 acts at inheriting generation to promote RNAi inheritance. *znfx-1(gg561)* animals expressing *pie-1∷gfp∷h2b* transgene (Ashe et al. 2012) were exposed to *gfp* dsRNA. A WT copy of *znfx1* was introduced by mating to WT males (or animals were allowed to self). Micrographs of GFP fluorescence in progeny oocytes are shown. In order to identify cross progeny the following strategy was employed. *znfx-1(gg561)* was marked by *dpy-10 (cn64) (dpy-10* is ~1 cM from *znfx-1). dpy-10 (cn64) 1+* animals are dumpy roller and *dpy-10 (cn64)* homozygous animals are dumpy, *znfx-1* genotypes was inferred based upon roller and dumpy phenotypes.

**Extended Data figure 7.**
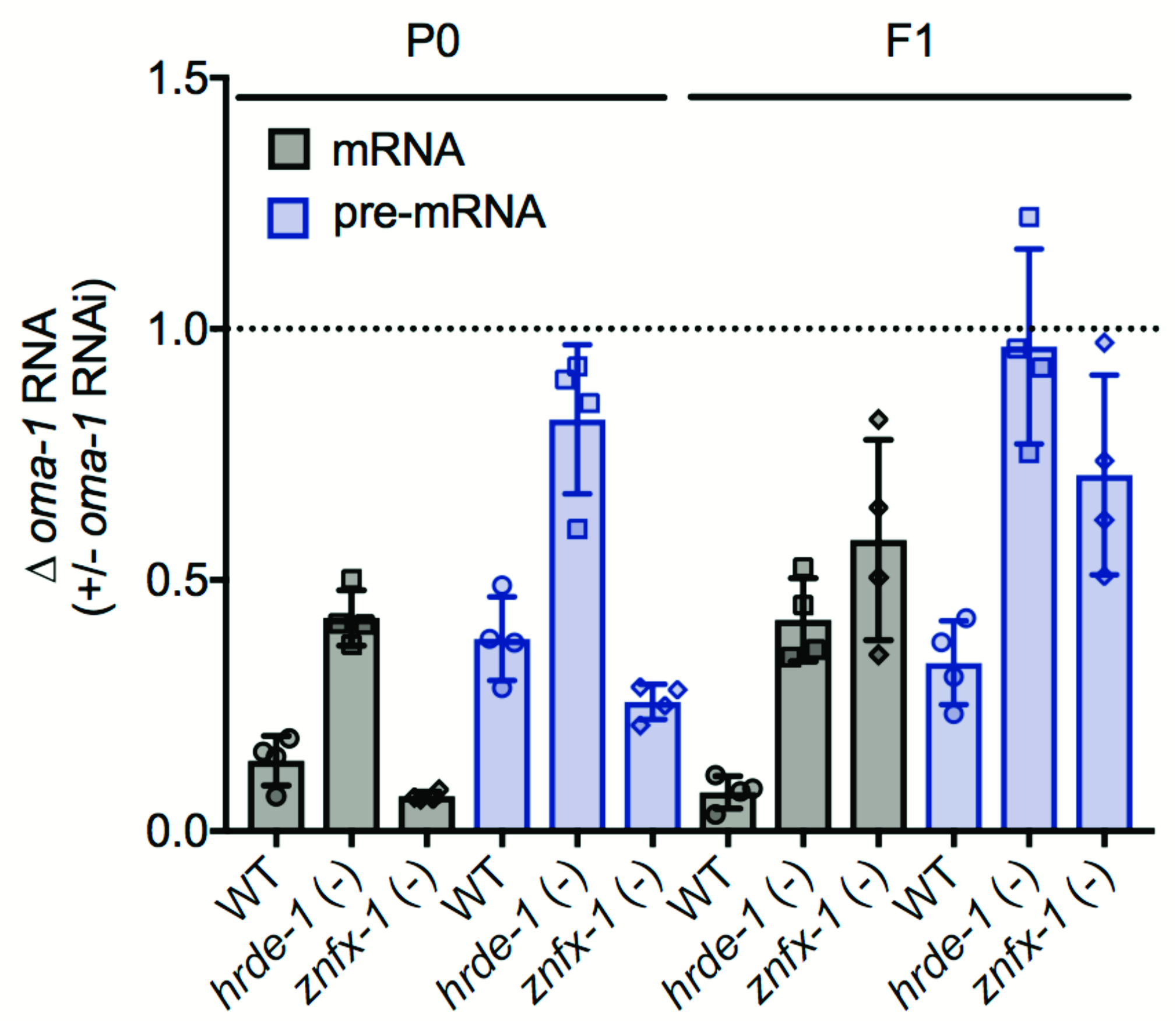
An independent primer set gives results similar to what we show in Figure 1 of main text. WT, *hrde-1(tm1200)*, and *znfx-1(gg561)* animals were exposed to *oma-1* dsRNA. Total RNA from RNAi (P0) and inheriting (F1) generations was isolated. *oma-1* RNA levels were quantified using qRT-PCR using primers 5’ to the site of RNAi and data were normalized to *eft-3* pre-mRNA level. n=4. Error bars are +/− standard deviation of the mean (SD).

**Extended Data Figure 8.**
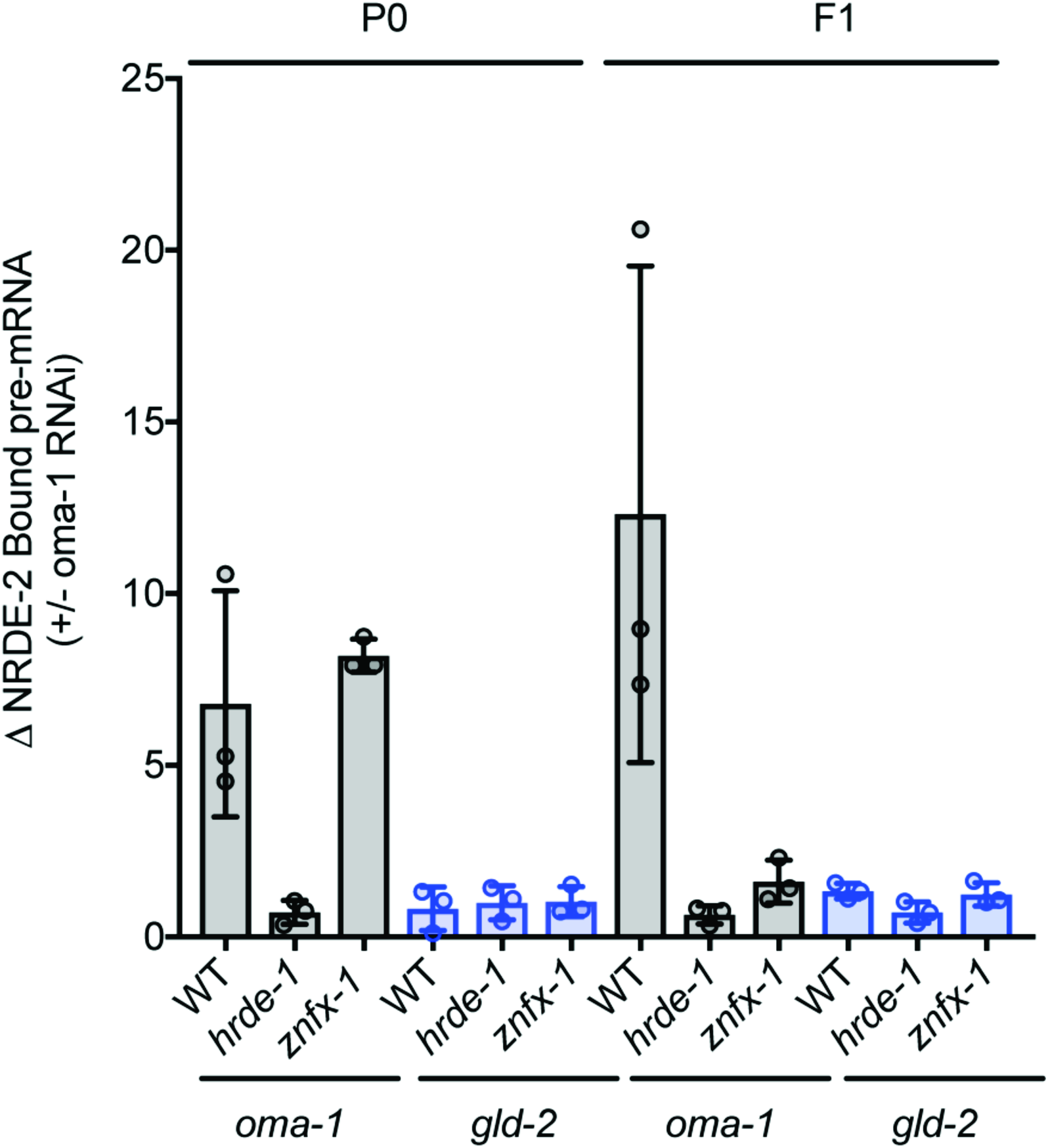
*znfx-1* is required for NRDE-2 to bind pre-mRNA only during inheritance phase of RNAi inheritance. Animals expressing NRDE-2∷3xFI_AG were treated with +/− *oma-1* RNAi. Extracts were generated from these animals as well as the progeny of these animals (which were not treated directly with *oma-1* RNAi). NRDE-2∷3xFLAG was IP’ed with an anti-FLAG antibody and NRDE-2 co-precipitating *oma-1* pre-mRNA was quantified by qRT-PCR with exon/intron primer sets designed to detect unspliced RNAs (pre-mRNAs) of the *oma-1* gene as well as a control germline expressed pre-mRNA *gld-2.* Data are expressed as ratio of signals ± *oma-1* RNAi. (n=3, bars, S.D).

**Extended Data Figure 9.**
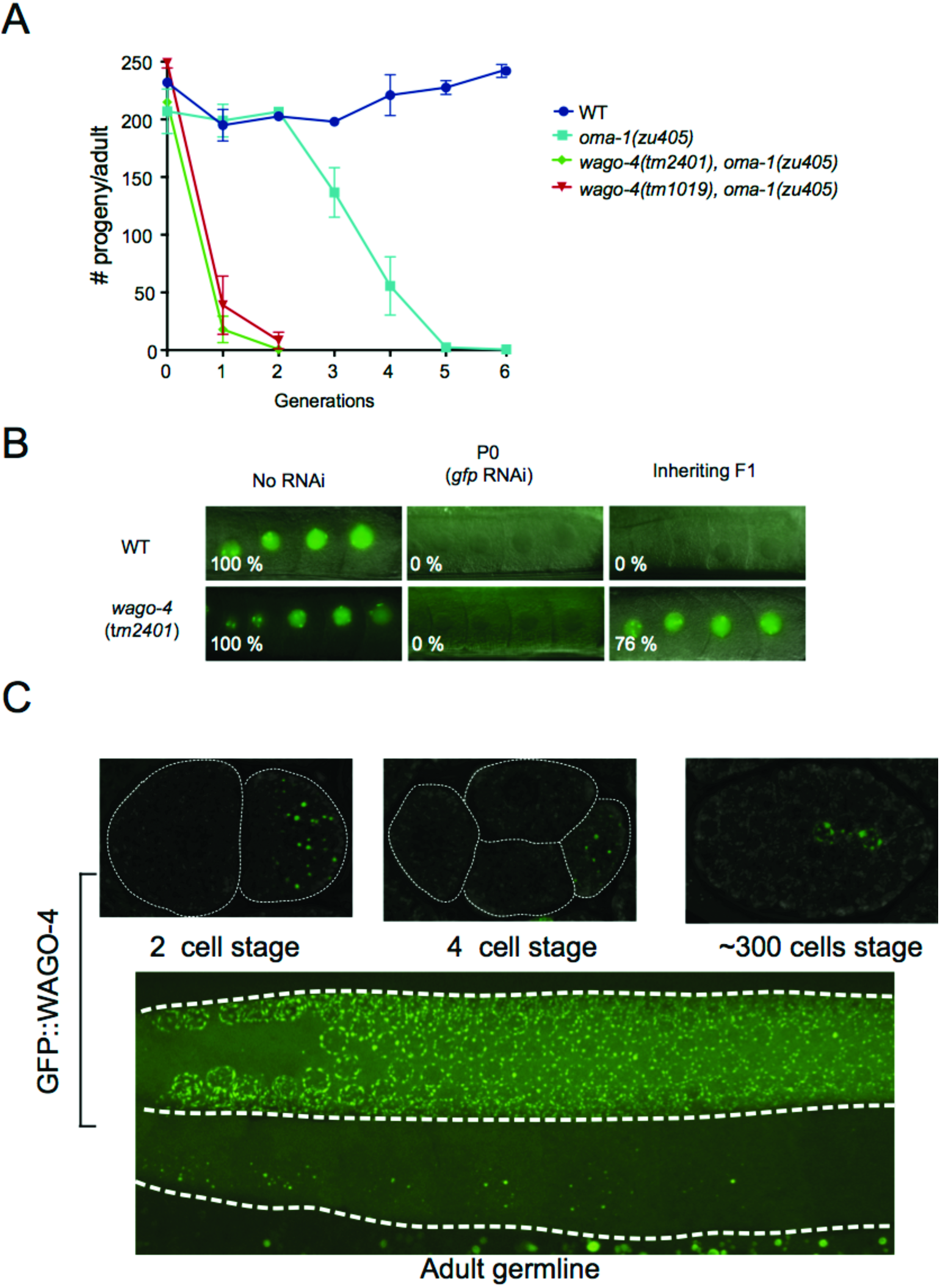
WAGO-4 is an Argonaute that localizes to the peri-nucleus and is required for RNAi inheritance. **A)** *oma-1(zu405)* is a temperature sensitive lethal (embryonic arrest at 20C) allele of *oma-1. oma-1* RNAi suppresses *oma-1(zu405)* lethality and this effect is heritable (Alcazar, Lin, and Fire 2008). Animals of indicated genotypes were exposed to *oma-1* dsRNA and F1 to F5 progeny were grown in the absence of *oma-1* dsRNA. Number of viable progeny of P0 (directly exposed to *oma-1* RNAi) and inheriting generations (F1 to F6, grown in the absence of *oma-1* RNAi) were scored (20°C). (n=3, bars, S.D). **B)** Animals of the indicated genotypes and expressing a *pie-1∷gfp∷h2b* transgene were exposed to *gfp* dsRNA (Ashe et al. 2012). Micrographs of animals +/− *gfp* RNAi as well as the F1 progeny of these animals are shown. % of animals expressing GFP is indicated. Percentages represent scoring of at least 90 animals in each generation and for each genotype. **C)** We used CRISPR/Cas9 to append a *gfp* tag upstream of the predicted *wago-4 atg* start codon. Top panels, fluorescent micrographs of *gfp∷wago-4* in 2 cell, 4 cell, and ~300 cell embryos. Bottom pane, fluorescent micrograph of the germline of an adult *gfp∷wago-4* animal.

**Extended Data Figure 10.**
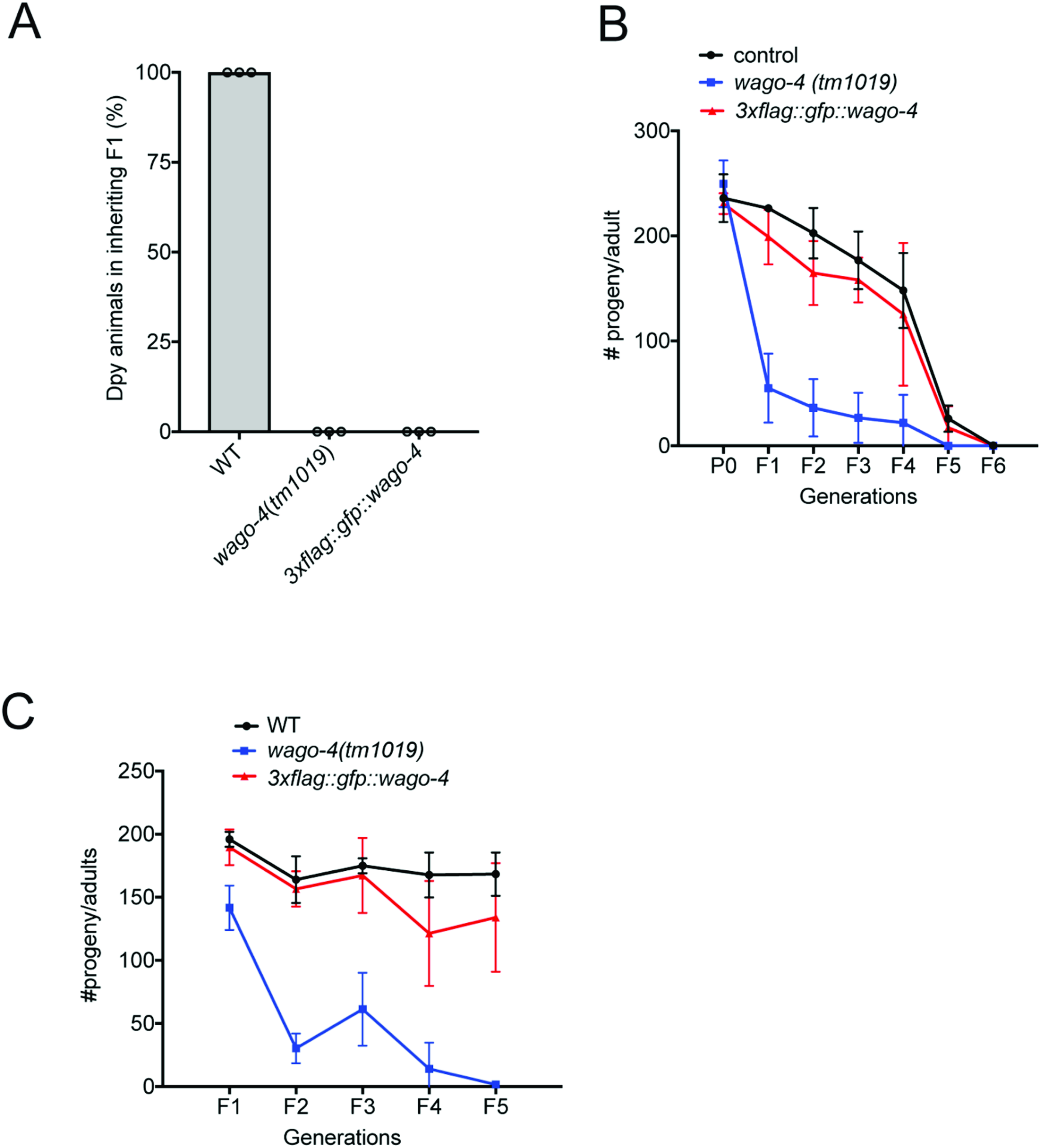
*3xflag∷gfp∷wago-4* encodes a partially functional protein. **A)** Animals of indicated genotypes were exposed to *dpy-11* dsRNA and F1 progeny were grown in the absence of *dpy-11* dsRNA. % of Dpy animals is shown. At least 150 animals of each genotype were scored. These data show that 3xFLAG∷GFP∷WAGO-4 is not functional for *dpy-11* inheritance. (n=3). **B)** This panel shows that 3xFLAG∷GFP∷ WAGO-4 is functional during *oma-1* RNAi inheritance. See Fig. S3B for details of *oma-1* RNAi inheritance assay. (n=3, bars, S.D). **C)** In figure 2D of the main text, we show that both *wago-4* and *znfx-1* exhibit mortal germline (Mrt) phenotypes at 25C. The data in this panel show that *3xflag∷gfp∷wago-4* animals are not Mrt, indicating that 3xFLAG∷GFP∷WAGO-4 is capable of promoting germline immortality. (n=3, bars, S.D).

**Extended data figure 11.**
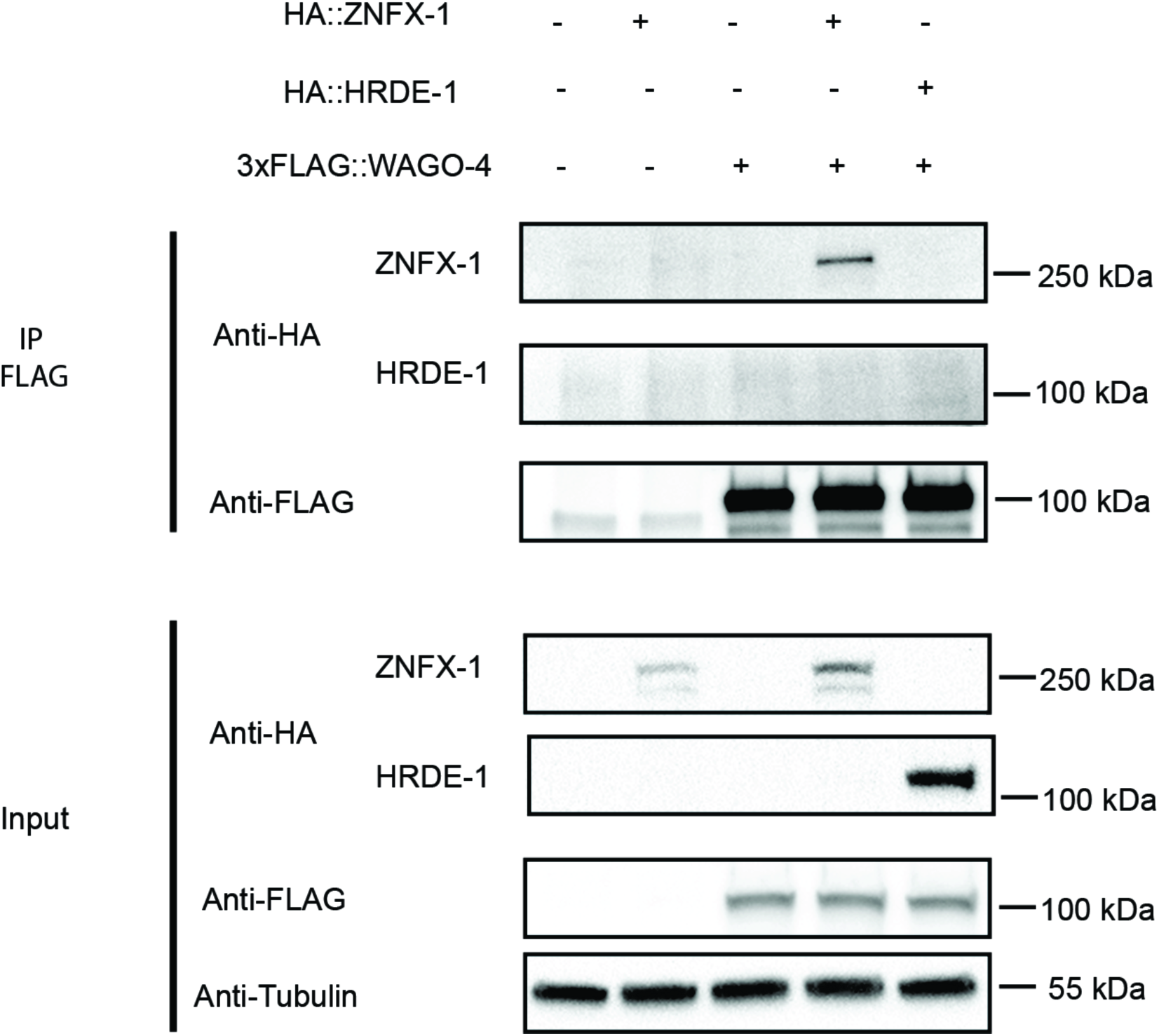
FLAG∷WAG0-4 does not co-IP with a control HA-tagged protein. CoIP analysis using animals expressing proteins with a *3xflag* tag appended to the *znfx-1 loci* or *ha* tags appended to the *wago-4* loci or the negative control *hrde-1* gene. KD (Kilodaltons). Data show that ZNFX-1 colPs with HA tagged WAGO-4, but not HRDE-1 control. HRDE-1 is an Argonaute protein that mediates TEI by directing TEI within nuclei (n=1).

**Extended Data Figure 12.**
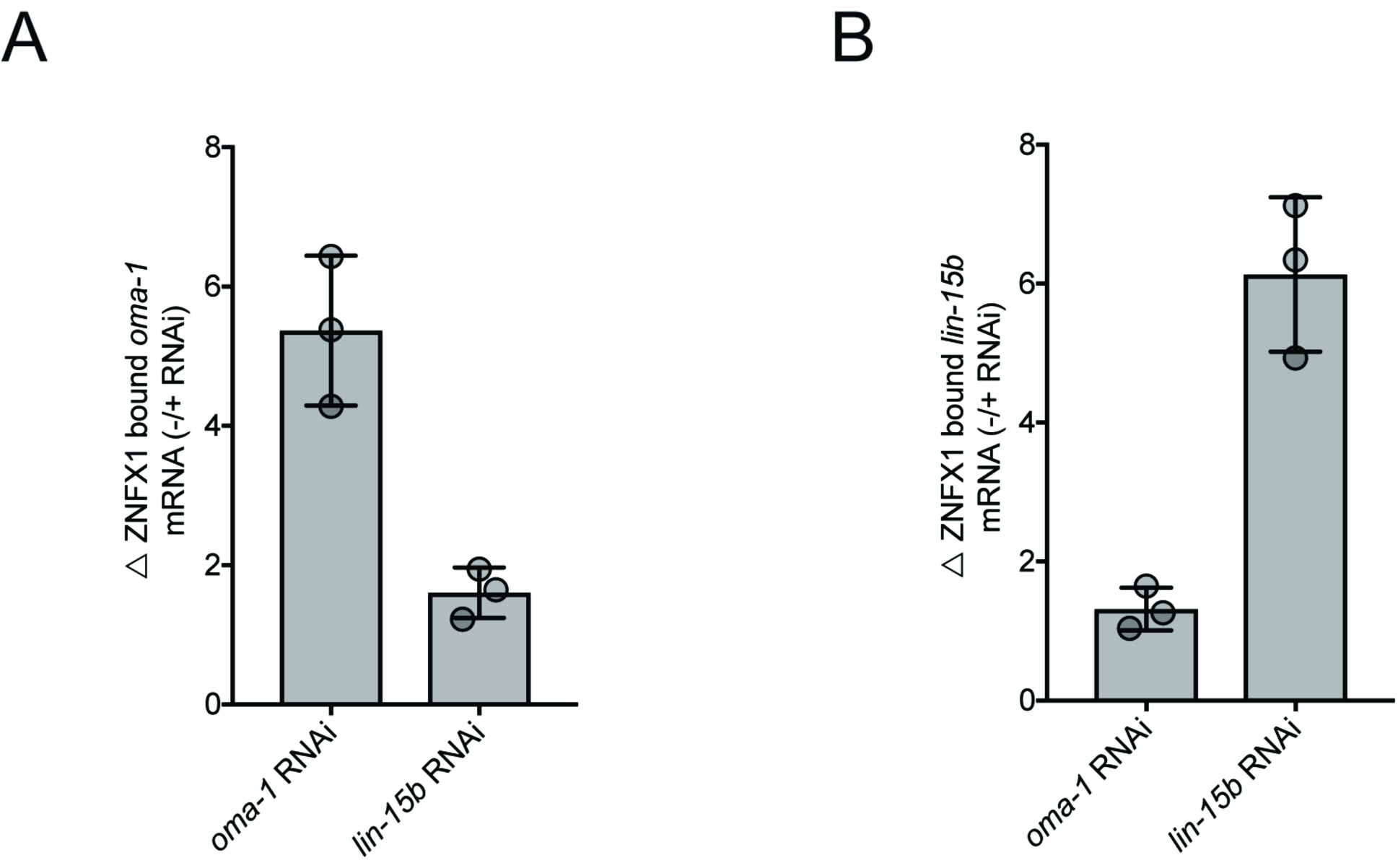
ZNFX-1/RNA interaction is sequence specific. Animals expressing 3xFLAG∷ ZNFX1 were treated with +/− *oma-1* RNAi or +/− *lin-15b* RNAi. Young adults were collected and lysed. 3xFLAG∷ZNFX1 was precipitated with an anti-FLAG antibody and **(A)** ZNFX1 co-precipitating *oma-1* mRNA or **(B)** *lin-15b* mRNA was quantified by qRT-PCR with mRNA specific primer set (one primer spanning two exons, another primer is in exon). Data are expressed as ratio ± RNAi. (n=3, bars, S.D).

**Extended Data figure 13.**
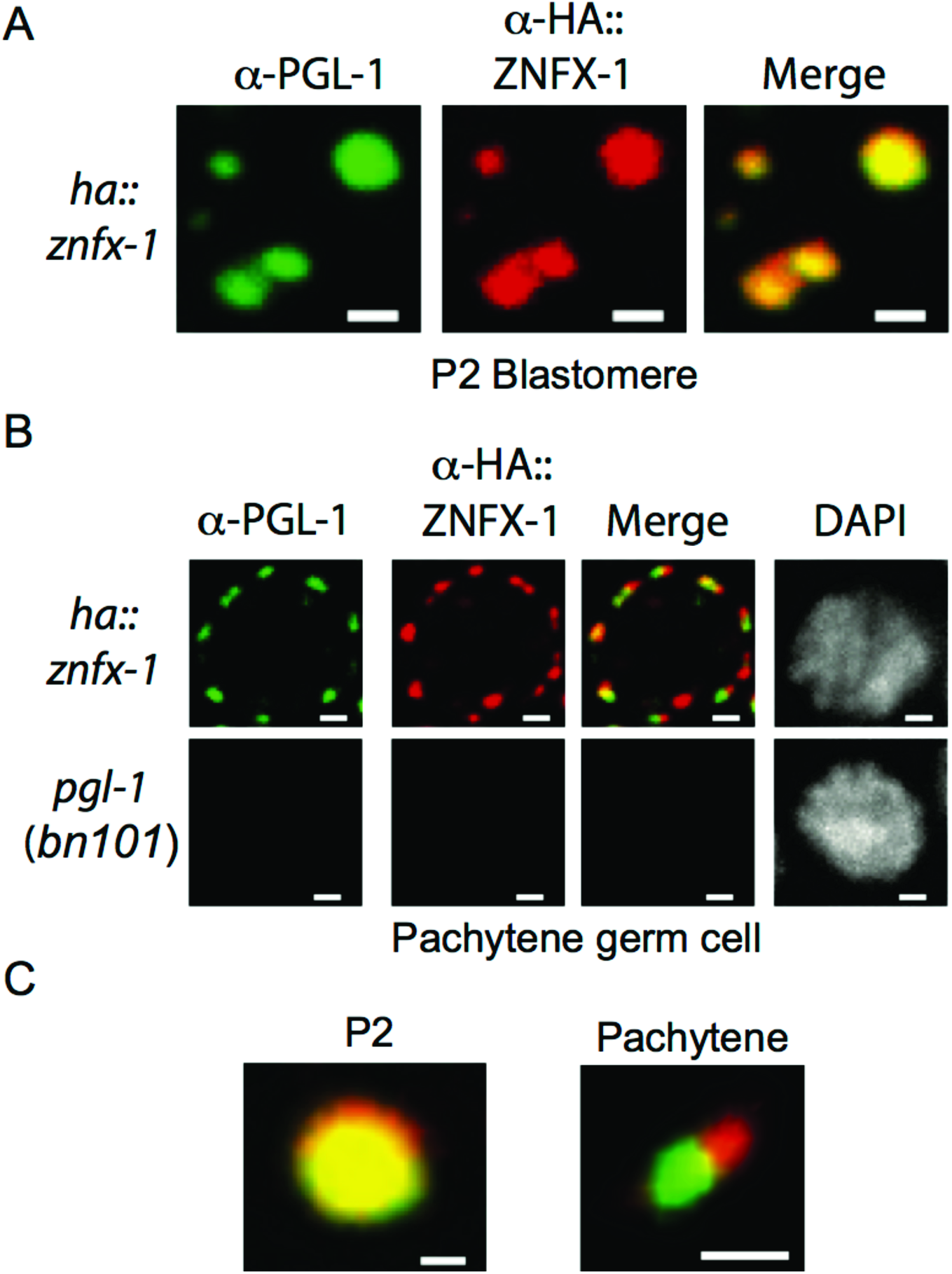
Visualization of Z granule formation with antibodies targeting PGL-1 (P granule) and HA∷ZNFX-1 (Z granule). To control for possible artifacts caused by fluorescent epitopes we conducted immunofluorescence on HA∷ZNFX-1 expressing animals using a-PGL-1 (K76 from Developmental Studies Hybridoma Bank) and a-HA (Abeam ab9110) antibodies. **A)** a-PGL-1 and a-HA signals colocalized in the P2 blastomeres of 4 cell embryos **B)**. α-PGL-1 and a-HA signals were adjacent, yet distinct in, in pachytene germ cells. No PGL-1 or HA∷ZNFX-1 signal was detected in *pgl-1(bn101)* animals, which do not express PGL-1 or HA∷ZNFX-1, establishing that IF signals were specific. **C)** Magnification of foci from A/B. Scale bars for **A)** = 1 μm, **B)** = 1 μm, **C)** = 0.5 μm.

**Extended Data figure 14.**
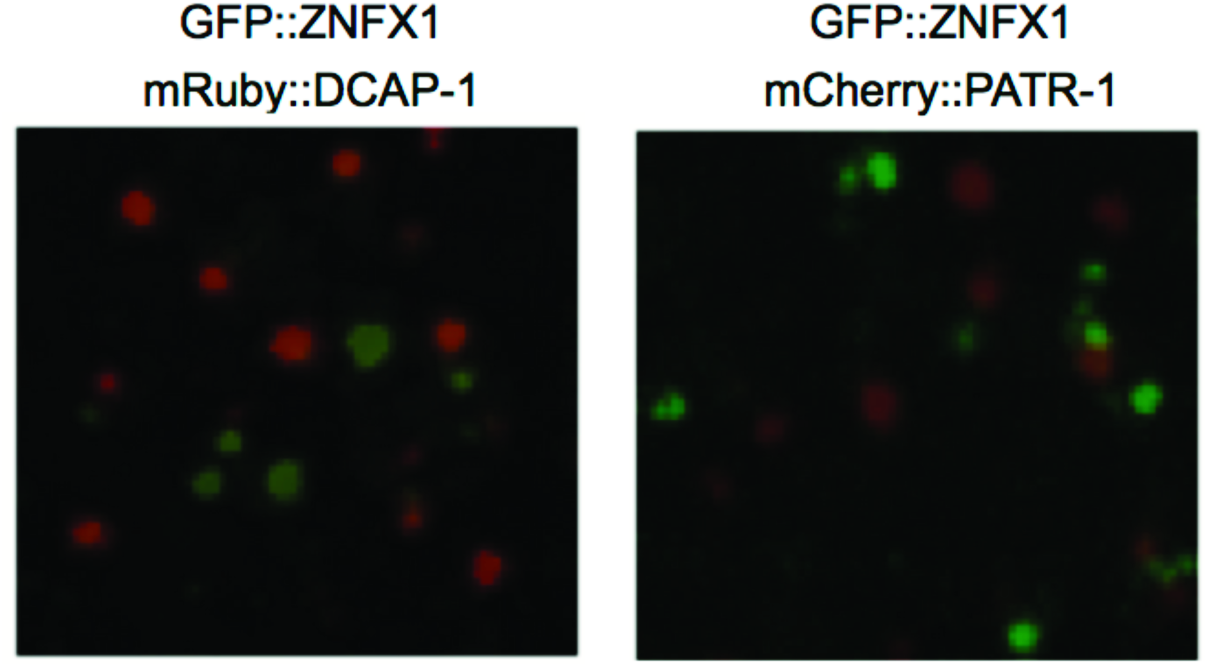
GFP∷ZNFX-1 does not colocalize with markers of processing bodies. PATR-1 and DCAP-1 localize to processing bodies (Gallo et al. 2008). Fluorescent micrographs of somatic blastomeres of embryos expressing the indicated fluorescent proteins. The data show that ZNFX-1 does not colocalize with markers of processing bodies in these cells.

**Extended Data figure 15.**
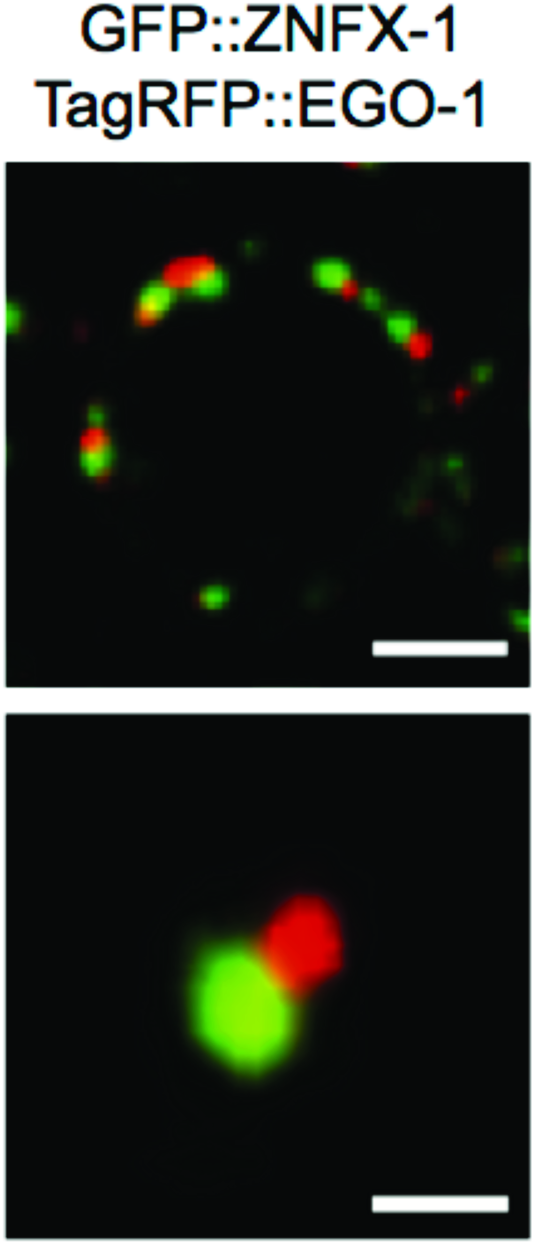
ZNFX-1 foci form adjacent to EGO-1 foci. Fluorescent micrographs of a single pachytene germ cell nucleus from animals expressing GFP∷ZNFX-1 and TagRFP∷EGO-1. 3D renders of a representative foci is shown below. Scale Bar = 0.5 μm

**Extended Data figure 16.**
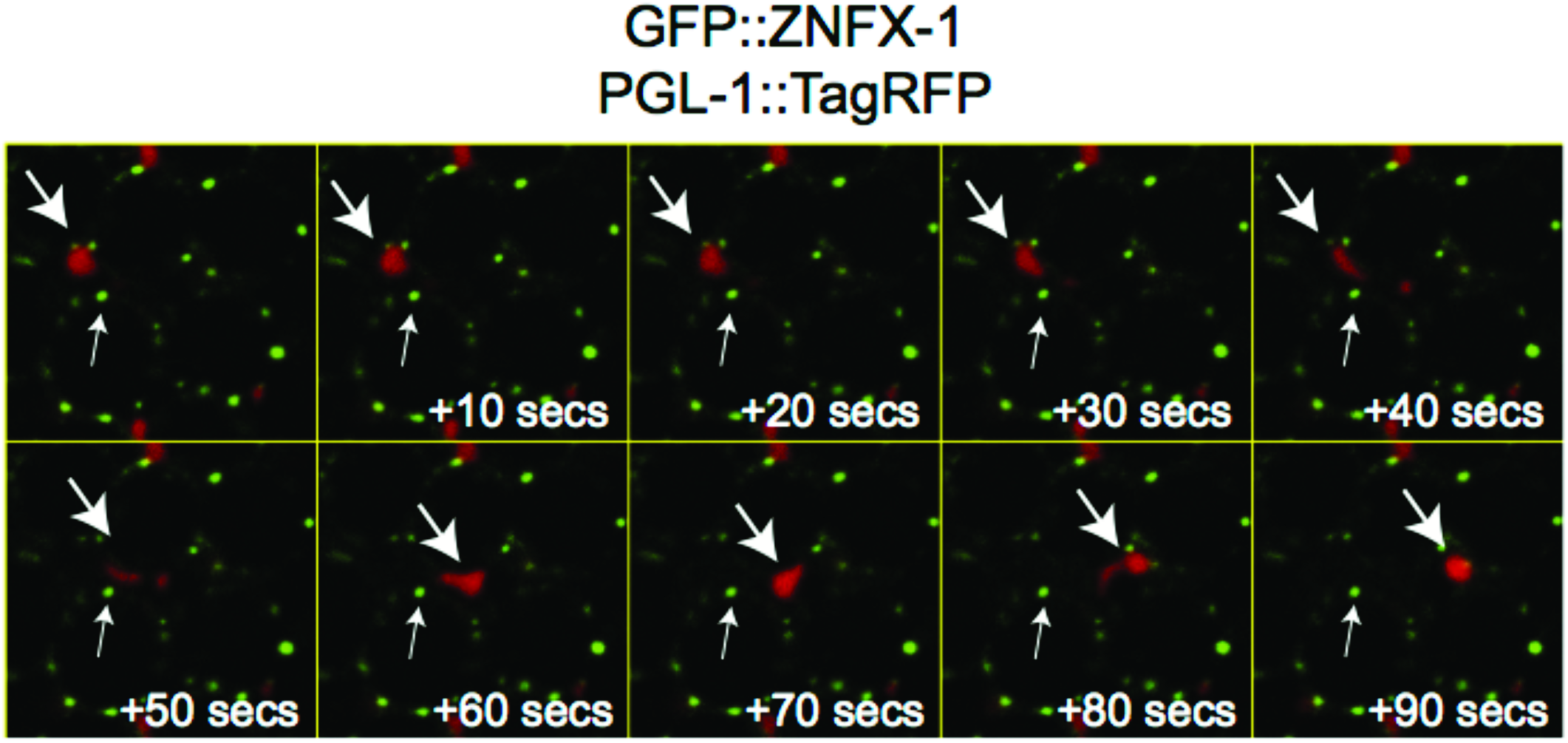
Z granules can be physically dissociated from P granules. Gonads were isolated from animals expressing GFP∷ZNFX-1 and PGL-1∷TagRFP and subjected to shearing force as described in (Brangwynne et al. 2009). Time lapse imaging at 10 second intervals is shown. A PGL-1-labeled P granule detaching from the nucleus and flowing throughout the cytoplasm is shown with (large arrow). ZNFX-1 labeled Z granules remain immobile (small arrow). Physical shearing was induced as previously described (Brangwynne et al. 2009). Briefly, GFP∷ZNFX-1 and PGL-1∷TAGRFP adults were dissected to extrude gonads. Isolated gonads were squeezed between two coverslips to generate shearing force. Coverslips were then mounted on a slide and imaged immediately with a spinning disc confocal. Z stacks were acquired every 10 seconds.

**Extended Data figure 17.**
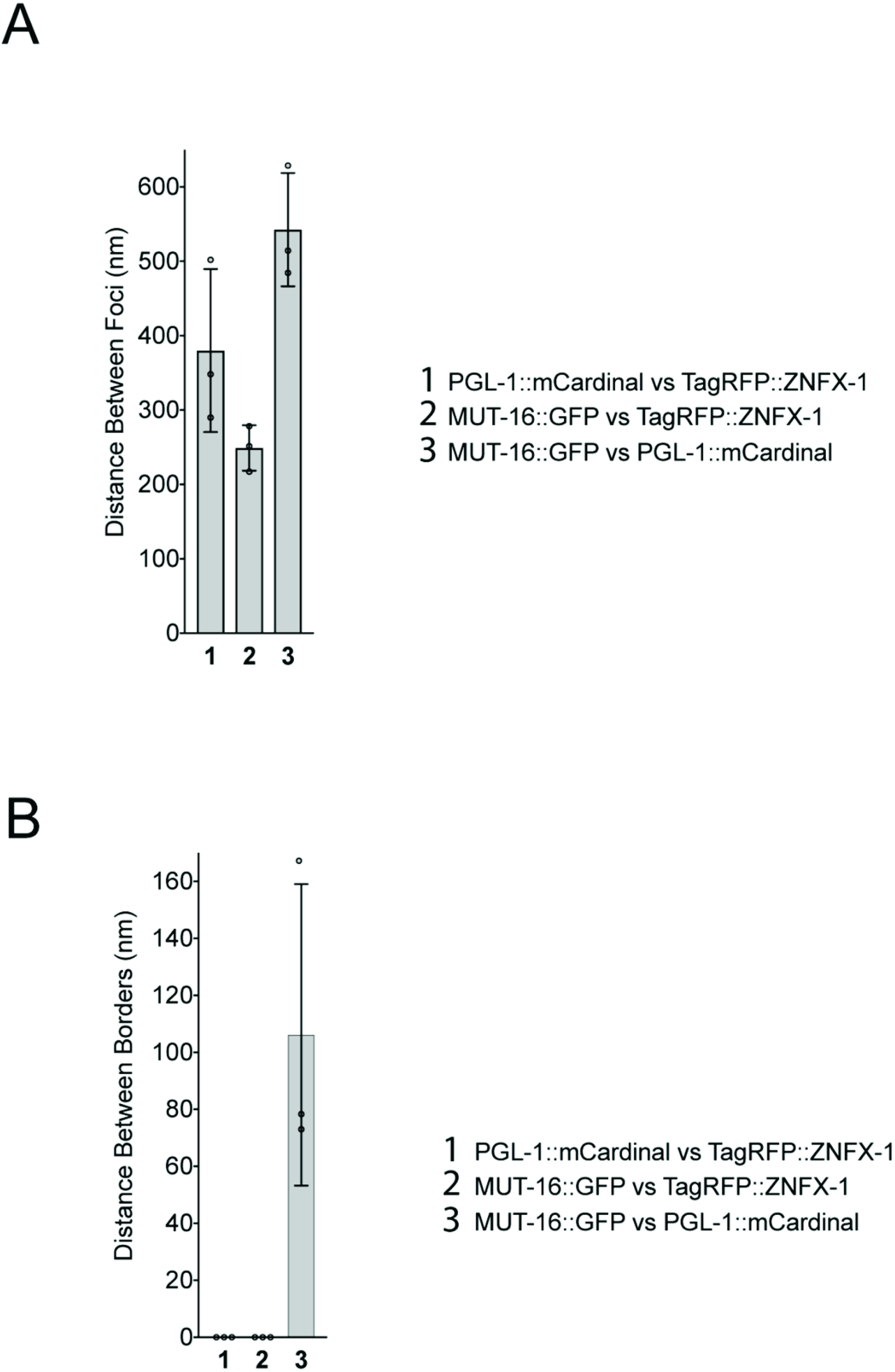
Quantification of centers and surfaces of fluorescence for Z granules, P granules, and *Mutator* foci. Distance between centers **(A)** and surfaces **(B)** of the spaces occupied by PGL∷mCardinal, tagRFP∷ZNFX-1, and MUT-16∷GFP were calculated as described in Methods. n= 3 (10 granules per sample) +/− SD.

**Extended Data figure 18.**
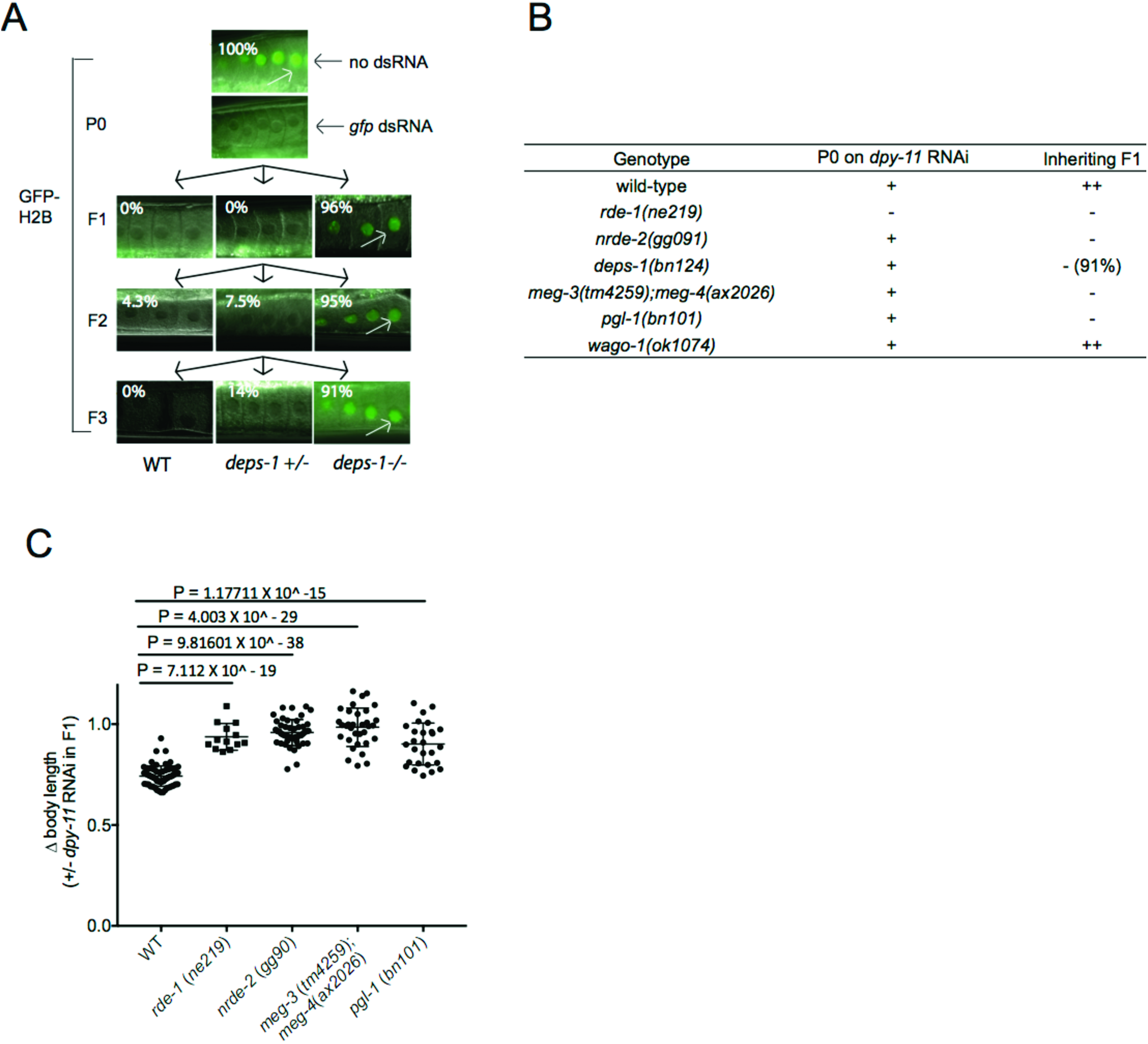
P granule assembly factors contribute to RNAi inheritance. **A)** DEPS-1 is required for P granule formation in adult germ cells ((Kawasaki et al. 1998; Spike et al. 2008; Wang et al. 2014). *deps-1 (bn124)/+)* animals expressing *pie-1∷gfp∷h2b* transgene (Ashe et al. 2012) were exposed to *gfp* dsRNA. Progeny were grown in absence of *gfp* dsRNA for three generations. Fluorescent micrographs show GFP expression in oocytes. % of animals expressing *pie-1∷gfp∷h2b* is shown. These data show that DEPS-1 activity is required in inheriting generations to allow for *gfp* RNAi inheritance. (n=32 for P0, n=100-110 for F1 to F3). **B)** MEG-3/4 and PGL-1 also contribute to P granule formation (Kawasaki et al. 1998; Spike et al. 2008; Wang et al. 2014). *dpy-11* RNAi causes animals exposed to *dpy-11* dsRNA to become Dumpy (Dpy). Progeny of animals exposed to *dpy-11* dsRNA inherit *dpy-11* silencing and are Dpy (Burton, Burkhart, and Kennedy 2011). RNAi inheritance mutants become Dpy in response to *dpy-11* RNAi however, progeny fail to inherit *dpy-11* silencing, and, therefore, are not Dpy. Animals of indicated genotypes were exposed to *dpy-11* dsRNA. F1 progeny were grown in the absence of *dpy-11* dsRNA. (-) indicates Non-Dpy, (+) indicates mild Dpy phenotype, (++) indicates strong Dpy (*pgl-1, deps-1*, and *meg-3/4)* are defective for *dpy-11* RNA inheritance. C) Animals of indicated genotypes were exposed to *dpy-11* dsRNA and F1 progeny were grown in the absence of *dpy-11* dsRNA. Body length of F1 animals were measured by Image J. Data are expressed as body length from progeny of *dpy-11* RNAi treated animals divided by average body length from control animals. (n=12-63. P values, student’s two tail ttest). These data are quantification of data presented in **(B)**.

**Extended Data figure 19.**
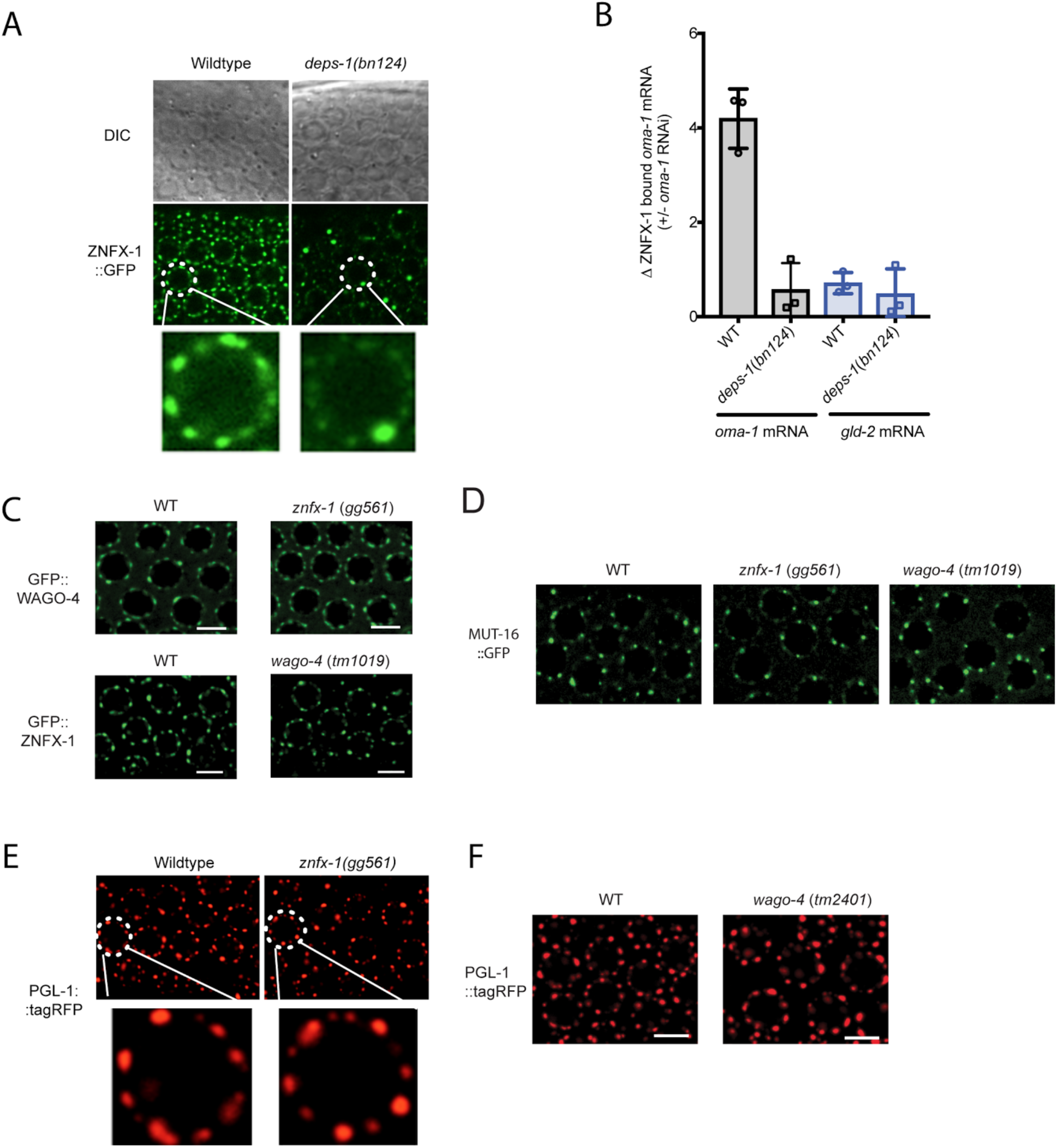
DEPS-1 is required for normal Z granule morphology and for ZNFX-1 to bind TEl-associated RNAs. **A)** In *deps-1(bn124)* animals, most Z granules are smaller than normal while typically one Z granule/nucleus becomes enlarged. Images are from pachytene region of germline. **B)** In *deps-1(bn124)* animals, ZNFX-1 doesn’t bind RNA. Wild-type or 3xFI_AG∷ZNFX-1 expressing animals were treated with *oma-1* dsRNA. ZNFX-1 was IP’ed in RNAi generation with a-FI_AG antibodies and co-precipitating RNA was subjected to qRT-PCR to quantify *oma-1* mRNA co-precipitating with ZNFX-1 in wild-type or deps-1(bn124) animals, *gld-2* is a germline expressed control mRNA. n=3. +/−SD. Panels **C-F** show that loss of ZNFX-1 or WAGO-4 does not seem to affect the formation of P granules **(E,F)** marked by PGL-1∷RFP, Z granules **(C)** marked by GFP∷WAGO-4/GFP∷ZNFX-1, or Mutator foci (D) marked by MUT-16∷GFP Note, in late embryonic germline development, PGL-tagRFP foci may not be efficiently concentrated into Z2/Z3 in *wago-4* mutant (data not shown).

**Extended Data figure 20.**
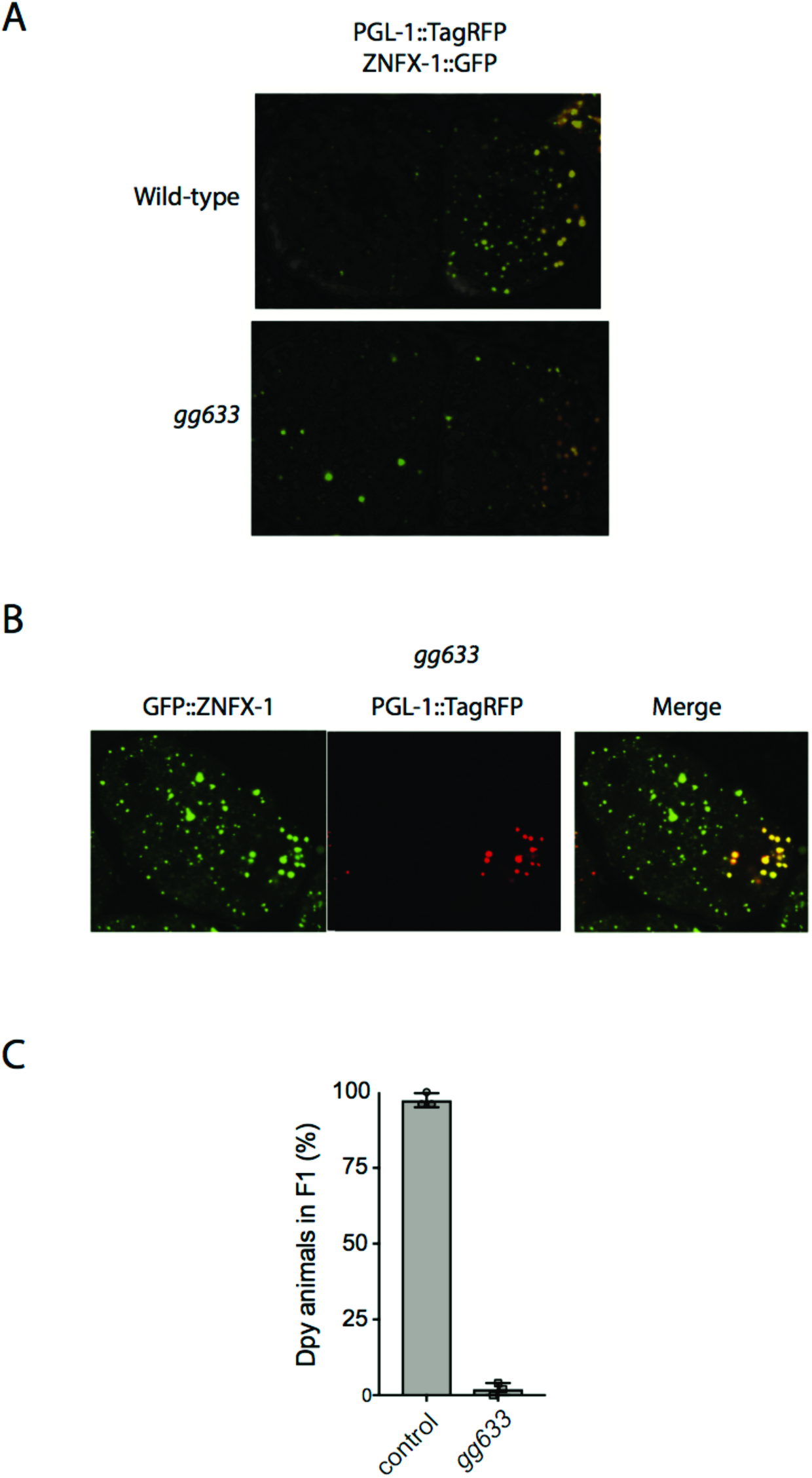
*gg633* is a mutation that inhibits the segregation of Z granules with the germ line as well as disrupts RNAi inheritance. **(A-B)** We conducted a pilot genetic screen to identify suppressors of *dpy-11* RNAi inheritance. We scored 26 *dpy-11* RNAi suppressors for Z granule morphology and Z granular germline segregation. In one of our mutant strains (*gg633)*, ZNFX-1 formed foci but these foci were not efficiently concentrated into the P lineage. **(A)** 2 cell embryo. **(B)** 4 cell embryo. The ZNFX-1 ∷GFP that did localize to P lineage appeared colocalized with PGL-1∷TagRFP **(C)** As expected, *gg633* animals are defective for *dpy-11* RNAi inheritance (n=3, +/− SD).

**Extended Data figure 21.**
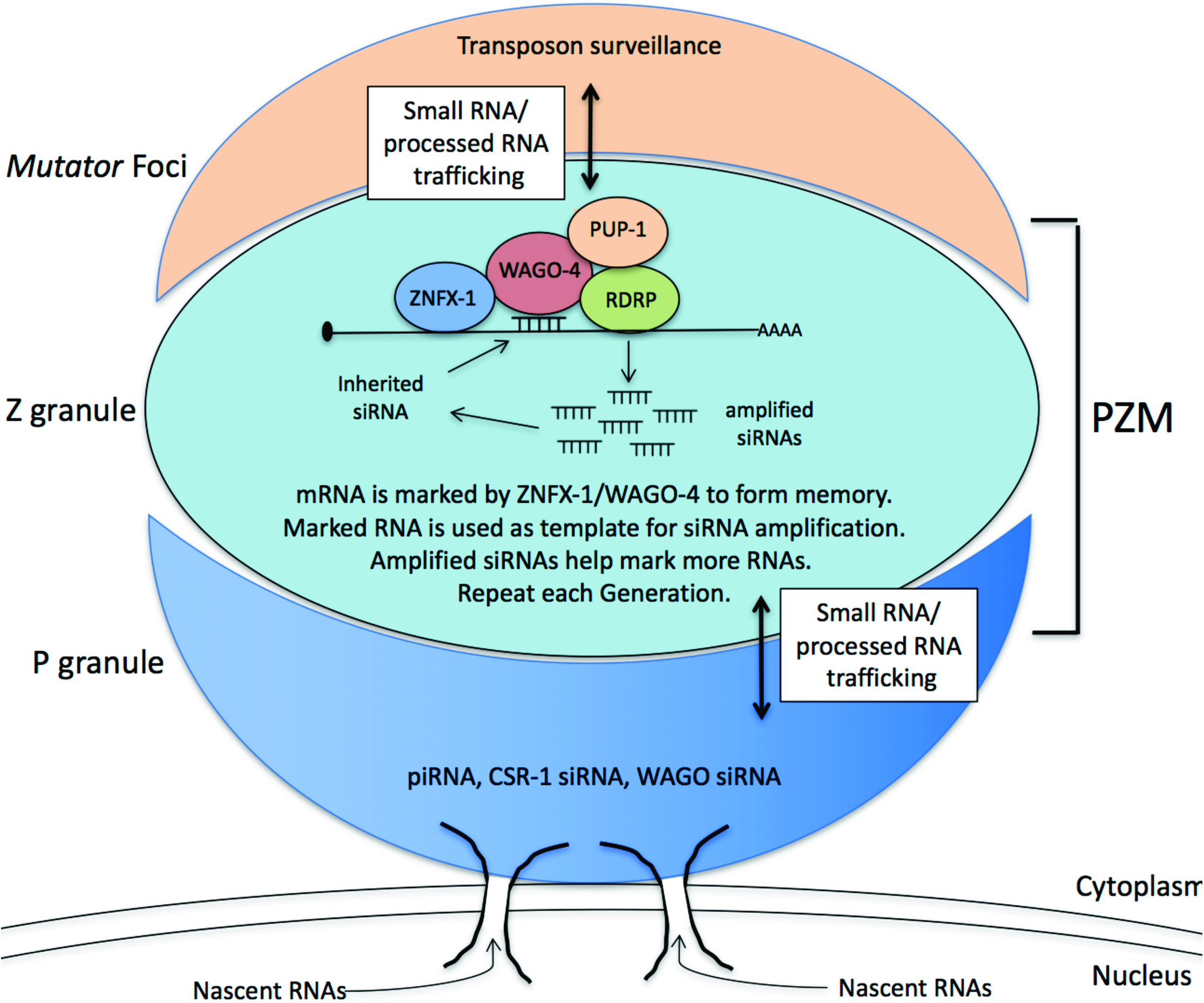
Working model for role of PZM assemblages in RNAi inheritance. P granules make contacts with nuclear pores (Pitt, Schisa, and Priess 2000; Sheth et al. 2010). The relationship between the Z and M segments of the PZM holo-droplet and nuclear pores are not yet known.

